# BTK dynamics and function are differentially influenced by distinct covalent inhibitor warhead chemistries

**DOI:** 10.64898/2026.05.07.723540

**Authors:** Raji E. Joseph, Robert G. Britton, David Yin-wei Lin, Julien Roche, Jeffrey A. Purslow, D. Bruce Fulton, Poowadon Fukasem, M. Paul Gleeson, Martin J. S. Dyer, Thomas E. Wales, Amy H. Andreotti

## Abstract

Covalent inhibitors are rapidly becoming the standard of care for treatment of a range of disease states. Covalent inhibitors bind irreversibly to their target using a reactive electrophile (or ‘warhead’). Acrylamide and 2-butynamide are the most commonly used cysteine targeting electrophiles. These warheads are chosen for their efficient and selective modification of the protein and are presumed to be otherwise functionally inert. Using a panel of BTK covalent inhibitors (Tirabrutinib, Acalabrutinib, Ibrutinib and Zanubrutinib), we show that the 2-butynamide warhead on Tirabrutinib and Acalabrutinib, unlike the acrylamide warhead on Ibrutinib and Zanubrutinib, induces conformational heterogeneity in key regions required for BTK signaling. Tirabrutinib or Acalabrutinib bound BTK adopt multiple conformational states that are in dynamic exchange, show increased binding to the substrate PLC_γ_ and are less effective at inhibiting PLC_γ_ signaling when compared to Ibrutinib. Swapping only the warheads between Tirabrutinib and Ibrutinib leads to a corresponding switch in BTK dynamics and inhibitor efficacy. The unanticipated warhead-specific allosteric effects raise interesting possibilities regarding inhibitor-specific mechanisms of resistance.

## INTRODUCTION

There are at least fifty covalent inhibitors approved for clinical use that target a wide range of proteins including viral proteases, G proteins and kinases ^1^. Covalent inhibitors are developed by attaching an electrophile (or warhead) to optimized high affinity reversible binders obtained from high throughput screens ^2,3^. Notably, the same warheads are routinely used to generate covalent inhibitors and so, many clinically approved covalent inhibitors share the same warhead chemistry ^4,5^. Acrylamide and 2-butynamide are the most commonly used cysteine modifier warheads ^4^. These electrophiles are chosen for their low reactivity which decreases toxicity arising from off-target, non-specific modifications ^5^.

The multi-domain, non-receptor kinase Bruton’s tyrosine kinase (BTK) ^6^ has emerged as an attractive target for covalent inhibitors. BTK is expressed primarily in cells of lymphoid or myeloid origin such as B cells, mast cells and macrophages where it functions downstream of immune receptors including the B cell receptor (BCR) and Fc receptor ^6-8^. BTK inhibitors are used in the treatment of multiple B cell malignancies and are being evaluated for the treatment of myeloid disorders such as rheumatoid arthritis and multiple sclerosis ^9-11^. Seven of the eight approved drugs that target BTK (Ibrutinib, Acalabrutinib, Zanubrutinib, Tirabrutinib, Rilzabrutinib, Remibrutinib and Orelabrutinib) are covalent inhibitors ^12^. All of these inhibitors have demonstrated efficacy and a pipeline of additional therapeutically valuable BTK inhibitors are at various stages of clinical development ^8,13^. Motivating this drug pipeline is the fact that acquired inhibitor-specific resistance mutations arise in clinical settings ^14^. As treatment options continue to develop it is important to understand how the chemical features of available inhibitors affect the target BTK protein and ensuing downstream responses.

Crystal structures of BTK/inhibitor complexes show a common binding mode: all utilize BTK C481 in the hinge region for covalent attachment, with the binding moiety extending into the kinase active site ‘back’ pocket (Figure 1a). Covalent attachment of the BTK inhibitors Tirabrutinib and Acalabrutinib is mediated via the 2-butynamide warhead whereas Ibrutinib and Zanubrutinib have an acrylamide warhead (Figure 1b,c ^4,15^). While these distinct warheads offer differences in reactivity ^16^, once the covalent bond is formed it is presumed that the warheads on different BTK inhibitors serve solely as attachment modules. Whether the warheads on BTK inhibitors have additional impacts on BTK has not been evaluated.

**Figure 1:**
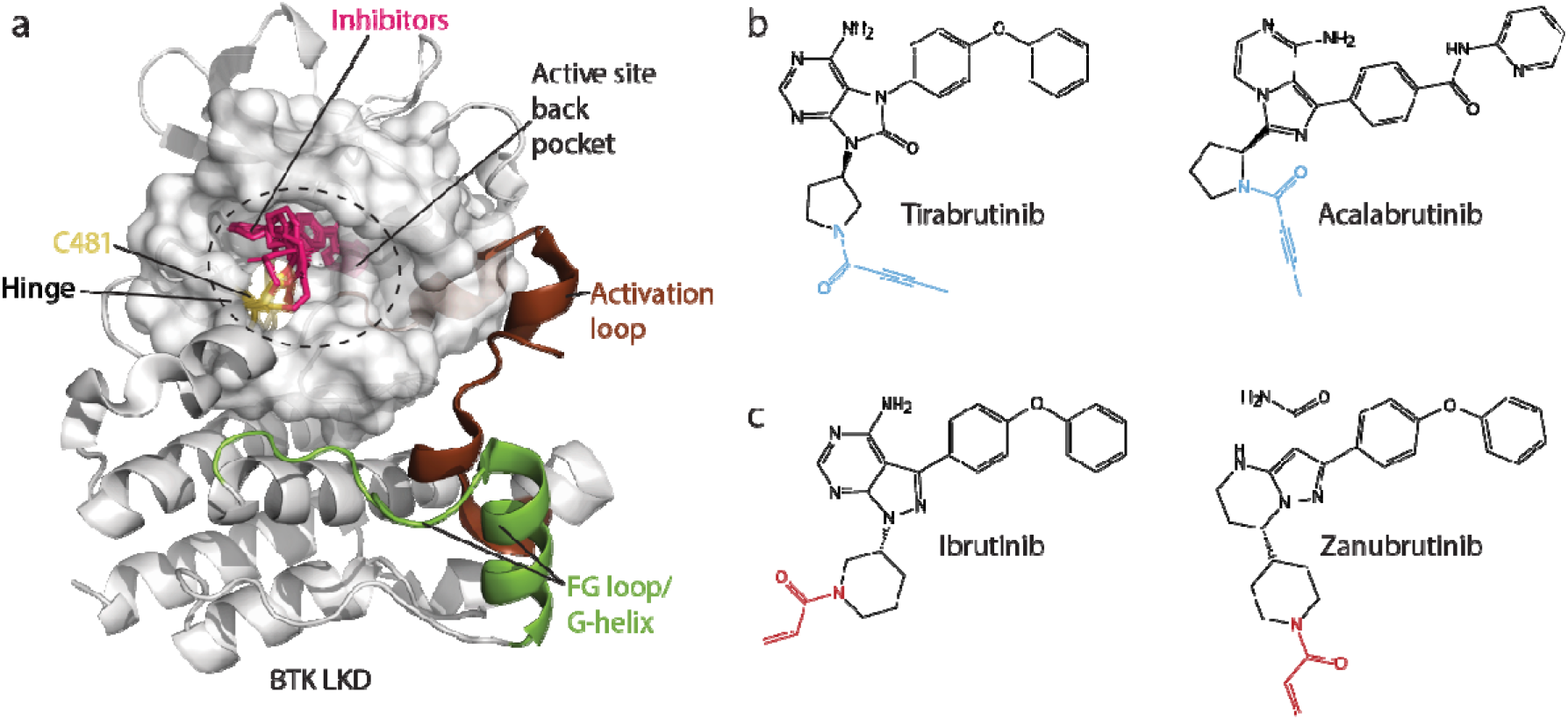
Covalent BTK inhibitors. (a) Structure of Ibrutinib bound BTK (PDB ID: 5P9J) overlaid with the structures of Tirabrutinib, Acalabrutinib and Zanubrutinib (pink). The kinase active site is shown as a space filling model. Covalent inhibitors bind to BTK C481 (yellow) and extend into the active site back pocket. The kinase activation loop (brown), hinge and the FG loop/G-helix (green) regions are labeled. (b,c) Structures of covalent BTK inhibitors used in this study. The electrophilic warheads are colored blue (2-butynamide) or red (acrylamide).

Our previous solution studies using Nuclear Magnetic Resonance (NMR) spectroscopy and Hydrogen-Deuterium Exchange-Mass Spectrometry (HDX-MS) have shown that a subset of BTK inhibitors exert unanticipated long-range effects on the conformation of full-length enzyme ^17,18^. Inhibitor characteristics that cause these structural changes and the functional consequences of inhibitor induced structural changes remain unexplored. Here we probe the determinants of specific inhibitor-induced allosteric changes in the BTK kinase domain using a combination of HDX-MS, NMR and computational analysis. We find that unique dynamic features induced by some inhibitors and not others have functional consequences. The observed conformational preferences are determined by the chemical nature of the covalent warhead. Inhibitors that utilize the 2-butynamide warhead, unlike the acrylamide warhead, induce multiple kinase domain conformational states resulting in increased deuterium uptake within the kinase domain, enhanced binding to the BTK substrate Phospholipase C_γ_ (PLC_γ_), and less efficient inhibition of PLC_γ_ signaling. Given the growing popularity of covalent inhibitors, the unanticipated allosteric effects driven by the nature of the covalent attachment should be considered in future inhibitor design and development.

## MATERIALS AND METHODS

### Constructs and reagents

All bacterial expression BTK and PLC_γ_1 cSH2 constructs have been described previously ^19-21^. Acalabrutinib, Zanubrutinib and Tirabrutinib were purchased from MedChem Express. Ibrutinib was purchased from Selleckchem. The acrylamide-Tirabrutinib and 2-butynamide-Ibrutinib analogs were synthesized as described in Supp. Data Files 1 and 2.

### Protein expression, purification and in vitro assays

Expression and purification of the PLC_γ_1 cSH2 and BTK kinase domain have been described previously ^19-21^. BTK/PLCγ interactions assay: 0.1 µM His-tagged BTK LKD was preincubated with no inhibitor (DMSO control) or 1.0 µM BTK inhibitor (Ibrutinib, Zanubrutinib, Acalabrutinib or Tirabrutinib) for 30 min at RT in pull-down buffer (20 mM Tris-HCl (pH 7.3), 150 mM Sodium chloride, 10% glycerol and 2% DMSO). 0.5 µM GST-tagged PLC_γ_1-cSH2 domain and 10 µL glutathione beads were added to Apo BTK LKD or the inhibitor bound BTK LKD and mixed for 2 hours at 4°C. The beads were washed thrice with the pull-down buffer and boiled in SDS-PAGE loading dye. The samples were separated by SDS-PAGE and western blotted with an Anti-His antibody (Millipore Sigma) ^19^. Western blots were developed by chemiluminescence and images captured using the ChemiDoc^TM^ gel imaging system (Biorad). The intensity of the bands was quantified using Image Lab (Biorad). The intensity of Ibrutinib bound BTK LKD was set to 1 and the intensity of the other bound BTK LKD is reported relative to Ibrutinib bound BTK. Data from three independent experiments were averaged and reported as mean ± SD.

### PLC_γ_2 phosphorylation assay

Ramos B cells (ATCC, # CRL-1596) were maintained in RPMI 1640 medium (Gibco, #A1049101) supplemented with 10% FBS (Thermo Fisher Scientific) and Penicillin/Streptomycin (Thermo Fisher Scientific) at 37°C/5% CO_2_. On the day of the assay, the cells were rinsed once with PBS and plated in RPMI 1640 medium without growth factors in a 60 mm dish at a density of 10 x 10^6^/ml cells and incubated with (50 nM final concentration) or without (0.1% DMSO final concentration) inhibitors for one hour at 37°C/5% CO_2_. The cells were harvested and kept on ice before stimulation. The cells were incubated for 10 min at 37°C and then stimulated with 20 µg/ml of F(ab’)_2_ anti-human IgM (Southern Biotech, # 2022-01) for varying time points (0, 1, 2 and 5 min) at 37°C. The samples were lysed in RIPA buffer containing protease inhibitors, sonicated briefly and then spun at 14K/10 min at 4°C. The supernatant was boiled with SDS-PAGE dye and analyzed by western immunoblotting for PLC_γ_2-pY759 (Cell Signaling, # 50535S) and PLC_γ_2 (Cell Signaling, # 34264S) as described earlier. The intensity of the bands was quantified using Image Lab (Biorad). The intensity of phosphorylated PLC_γ_2 in each time point is reported relative to the unstimulated sample. Data from three independent experiments were averaged and reported as median with interquartile range (IQR).

### Calcium flux assay and IC_50_ determination

Ramos B cells (ATCC, # CRL-1596) were maintained in RPMI 1640 medium (Gibco, #A1049101) supplemented with 10% FBS (Thermo Fisher Scientific) and Penicillin/Streptomycin (Thermo Fisher Scientific) at 37°C/5% CO_2_. One day prior to the calcium assay, the cells were rinsed once with PBS and plated in RPMI 1640 medium containing 1% FBS in 96 well clear bottom black plates (Corning, CellBIND) at a density of 100,000 cells/well and incubated for 18 h at 37°C/5% CO_2_. Inhibitors (0.1% DMSO final concentration) and the FLIPR calcium 6 assay reagent (Molecular Devices) were added to the cells for one hour at 37°C/5% CO_2_. Cells were stimulated with 5 µg/ml of F(ab’)_2_ anti-human IgM (Southern Biotech, # 2022-01) and fluorescence was monitored at 37°C for 6 minutes (Ex. 480 nm, Em. 545 nm) on a Tecan Spark plate reader. Percent inhibition of calcium flux was calculated using the no inhibitor control as the maximum. IC_50_ values for calcium flux inhibition were calculated using the AAT Bioquest IC_50_ calculator ^22^. IC_50_ values determined from six independent runs were averaged to determine the mean IC_50_ value. Each IC_50_ value was converted to pIC_50_ and averaged to determine the mean pIC_50_ ± standard deviation.

### NMR spectroscopy

Uniformly ^15^N labeled BTK samples were produced as described previously ^19^. All NMR spectra were acquired at 298 K on a Bruker AVIII HD 800 MHz spectrometer equipped with a 5 mm TCI (H/F)CN z-shielded gradient triple resonance cryoprobe, operating at a ¹H frequency of 800.37 MHz. NMR samples with inhibitors consisted of 150 µM ^15^N labeled BTK, mixed with 200 µM inhibitor and 2% DMSO unless indicated otherwise. All data were analyzed using NMRViewJ ^23^. Methods for high pressure NMR and ZZ-exchange experiments (and x-ray crystallography) are provided in Supplemental Data File 1.

### HDX-MS

Procedures for HDX-MS of BTK have been described previously ^17,18^. Experimental parameters are provided in the Supp. Data File 3 ^24^. All HDX-MS data have been deposited to the ProteomeXchange Consortium via the PRIDE ^25^ partner repository with the dataset identifier PXD069107. Comparisons have been made to previously published HDX MS data (taken from reference ^17^, identified with an asterisk in Supp. Data File 3 and a dataset identifier PXD047865 in the PRIDE partner repository). Deuterium incorporation was determined using Waters DynamX 3.0. Vertical difference maps in Figures 2,4, S1 and S4 do not represent a linear sequence of non-overlapping peptides. All coincident and overlapping peptides for comparisons in each figure are provided in figure identified tabs of the Supp. Data File 3. Intact mass analysis for 2-butynamide Ibrutinib and acrylamide-Tirabrutinib analogs bound to full-length (FL) BTK shows a single inhibitor bound per BTK FL (Supp. Fig. S8). Intact mass analysis for Ibrutinib, Zanubrutinib, Acalabrutinib or Tirabrutinib bound FL BTK has been previously published and also shows a 1:1 binding stoichiometry^17^.

**Figure 2:**
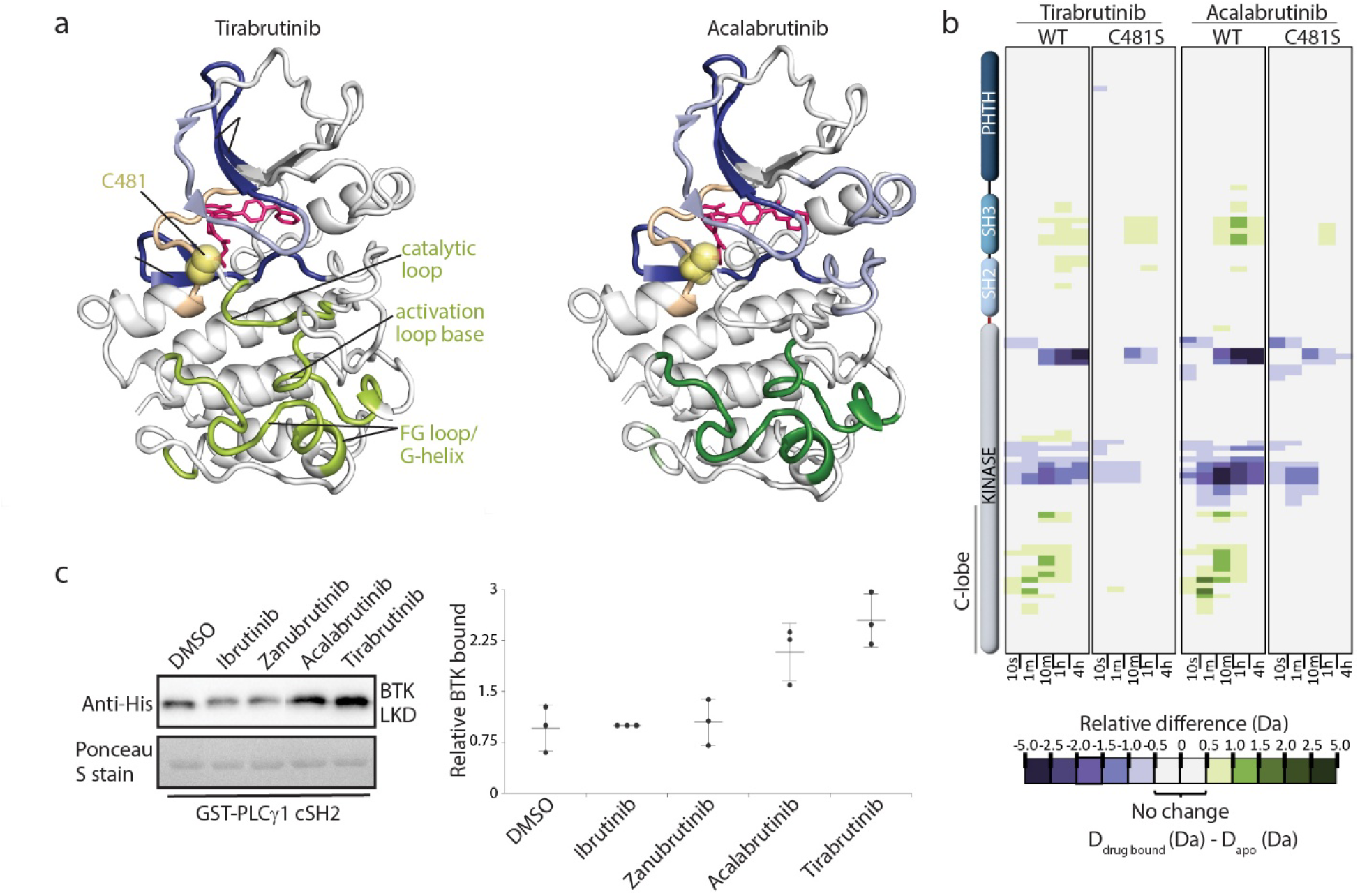
Dynamic allostery and in vitro functional effects of the distinct warheads. (a) Mapping the HDX-MS changes induced by inhibitor binding onto the structure of the BTK kinase domain. Differences > 1.0 Da are dark blue (decrease) or dark green (increase); differences 0.5 Da-1.0 Da are light blue (decrease) and light green (increase). Inhibitors are pink and C481 is yellow. Regions of increased deuterium uptake in the kinase domain C-lobe are labeled. (b) Covalent attachment of BTK with Tirabrutinib and Acalabrutinib is required to observe the dynamic changes in the C-lobe. Relative deuterium level of peptides in apo BTK is subtracted from the deuterium level of the corresponding peptide from each drug-bound form (D_drug-bound_-D_apo_); scale shows magnitude of differences. Peptic peptides are shown from BTK N- to C-terminus, top to bottom, and sample time in deuterium is shown left to right. (c) Tirabrutinib or Acalabrutinib bound BTK linker-kinase domain (LKD) shows increased binding to PLC_γ_ as compared to Ibrutinib and Zanubrutinib bound BTK LKD or Apo BTK LKD. Anti-His detects bound BTK, Ponceau stain shows total PLCγ cSH2. Bands were quantified and plotted as a dot plot (described in methods). Data from three independent experiments were averaged and reported as mean ± SD. Source data for this figure is in Supp. Data File 4.

### Statistics and Reproducibility

All experiments were carried out in triplicate to ensure reproducibility. Specific statistics applied to each experiment are detailed in either the figure caption or the methods section. Data are presented as mean with standard deviation or median with interquartile range.

## Data availability

All HDX-MS data have been deposited to the ProteomeXchange Consortium via the PRIDE ^25^ partner repository with the dataset identifier PXD069107. Comparisons have been made to previously published HDX MS data (taken from reference ^17^), has a dataset identifier PXD047865 in the PRIDE partner repository. The atomic coordinates and structure factors for the crystal structures have been deposited into the PDB with the accession codes: 9XYV (BTK KD/Ibrutinib), 9XYY (BTK KD/2-butynamide-Ibrutinib), and 9XZ2 (BTK KD/acrylamide-Tirabrutinib). All raw data underlying the figures are provided in Supp. Data Files 3 and 4.

## RESULTS

### Tirabrutinib and Acalabrutinib induce functionally important long-range conformational effects in the BTK kinase domain

HDX-MS analysis of covalent BTK inhibitors Tirabrutinib, Acalabrutinib, Ibrutinib, and Zanubrutinib have shown that drug binding to full-length BTK induces an increase in deuterium uptake in the SH3 and SH2 domains and a decrease in the kinase N-lobe surrounding the active site (Figure 2a,b, Supp. Fig. S1a,b ^17^). HDX-MS data also reveal differences across this panel of inhibitors; binding of Tirabrutinib or Acalabrutinib to BTK (but not Ibrutinib or Zanubrutinib) results in an increase in deuterium uptake in three regions of the kinase domain C-lobe, the catalytic loop, the base of the activation loop, and the FG loop/G-helix (Figure 2a,b Supp. Fig. S1a,b). Differences in the C-lobe among the panel of BTK/inhibitor complexes are not evident in crystal structures of the BTK/drug complexes (Supp. Fig. S2a-e).

The inhibitor specific dynamic differences in the FG loop/G-helix region of the C-lobe is of particular interest, as this region has been previously identified as a docking site required for BTK mediated phosphorylation of its substrate PLC_γ_ ^26,27^. To test whether the inhibitor induced conformational/dynamic changes in the FG loop/G-helix might affect substrate binding, we compared the interaction of PLC_γ_ with the different covalent inhibitor/BTK complexes. Apo His-tagged BTK linker-kinase domain (LKD) or BTK LKD bound to Ibrutinib, Zanubrutinib, Acalabrutinib or Tirabrutinib was incubated with GST-tagged PLC_γ_1 cSH2 (minimal fragment required for direct PLC_γ_/BTK interaction). BTK bound to Tirabrutinib or Acalabrutinib interacts more effectively with PLC_γ_ compared to apo BTK, Ibrutinib or Zanubrutinib bound BTK (Figure 2c). This finding suggests that the conformational/dynamic features unique to the FG loop/G-helix in the BTK/Tirabrutinib or BTK/Acalabrutinib complexes (Figure 2a,b) directly affect PLC_γ_ substrate interaction.

The observed differences in PLCγ binding are interesting in the context of previous studies reporting lower efficacy (higher IC_50_) for Acalabrutinib compared to Ibrutinib in terminating downstream PLC_γ_ signaling in B cells ^28,29^. To first probe whether Tirabrutinib, which like Acalabrutinib contains the 2-butynamide warhead, is similarly less effective than Ibrutinib we measured PLC_γ_2 phosphorylation post BCR stimulation. As shown in Figure 3a,b, Ibrutinib is more effective at inhibiting PLC_γ_2 phosphorylation compared to Tirabrutinib. Additionally, we measured IC_50_ values for Calcium flux post BCR stimulation for both drugs. Following previous reports for Acalabrutinib, we find that Tirabrutinib is less effective (IC_50_=50.8 nM) at terminating calcium signaling post BCR stimulation compared to Ibrutinib (IC_50_=5.8 nM), raising the possibility that the 2-butynamide warhead containing BTK inhibitors might generally be less efficient at terminating PLC_γ_ signaling (Figure 3c). The difference in IC_50_ values we measure for Tirabrutinib versus Ibrutinib (approaching ten-fold) is on the same order of magnitude reported previously for Acalabrutinib and Ibrutinib ^30^.

**Figure 3:**
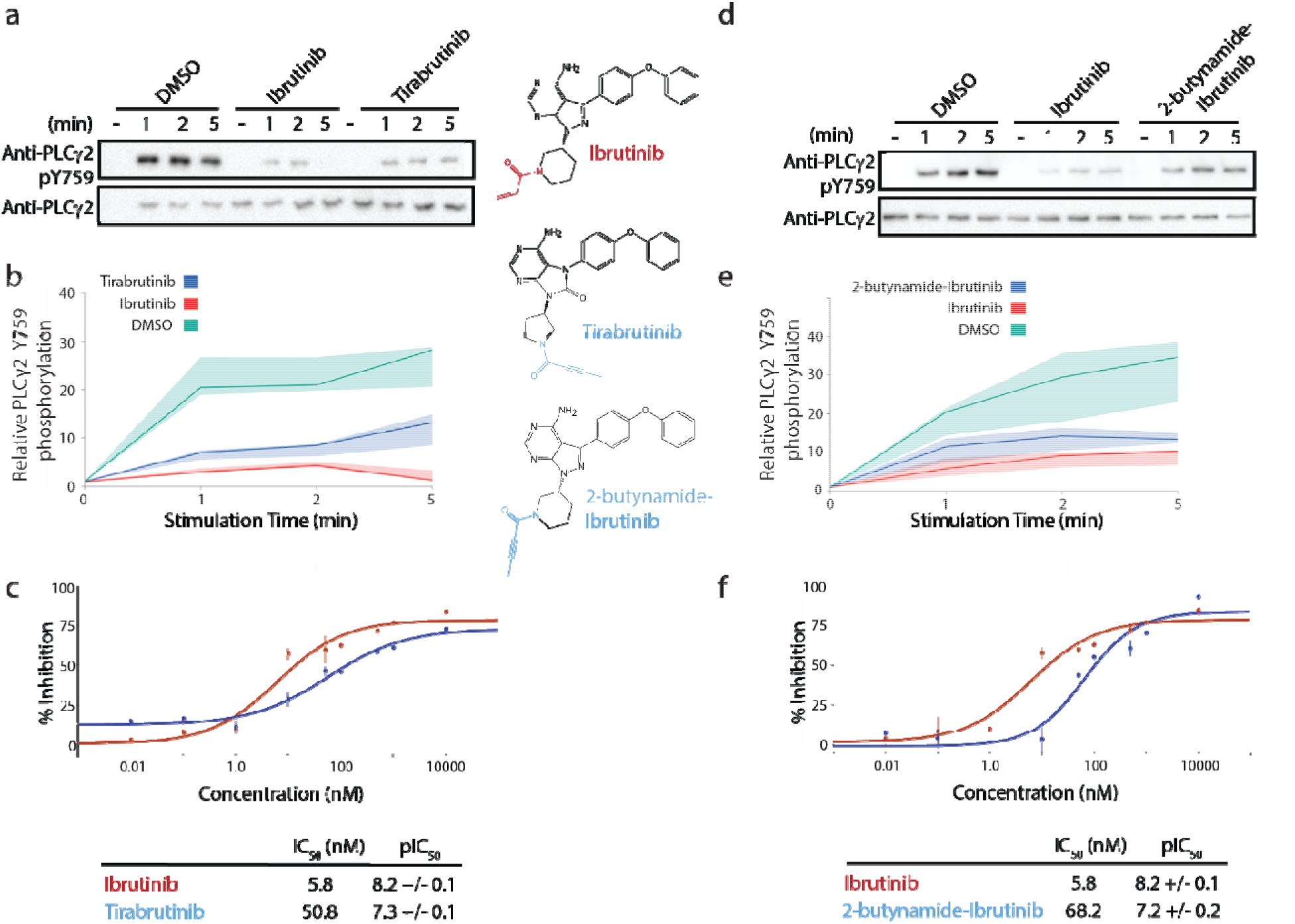
The acrylamide warhead containing BTK inhibitors are more effective at terminating downstream BCR signaling compared to 2-butynamide warhead containing BTK inhibitors. (a) Ibrutinib is more effective at inhibiting PLC_γ_2 phosphorylation in Ramos B cells post BCR stimulation compared to Tirabrutinib. Structures of inhibitors (and analog) are shown highlighting the 2-butynamide (blue) and acrylamide (red) warheads. (b) Quantification of the data shown in (a) with details in the methods. Data from three independent experiments were averaged and reported as median (bold lines) with interquartile ranges (IQR, shaded areas). (c) Ibrutinib (red) is more effective than Tirabrutinib (blue) at inhibiting calcium flux in BCR stimulated Ramos B cells. Half maximal inhibitory concentration (IC_50_) values and -log_10_ IC_50_ (pIC_50_) values ± SD (n=6) of calcium flux inhibition are listed. (d) The 2-butynamide-Ibrutinib analog with the Tirabrutinib warhead is less effective at inhibiting PLC_γ_2 phosphorylation a compared to Ibrutinib. (e) Quantification of the data shown in (d) was carried out as in (b). (f) IC_50_ for calcium flux inhibition for 2-butynamide-Ibrutinib (blue) is higher than that of Ibrutinib (red) and is similar to that of Tirabrutinib. Data are as described in (c). Source data for this figure is in Supp. Data File 4.

### The covalent attachment site is responsible for conformational dynamics in the kinase domain C-lobe

There are several chemical differences between Tirabrutinib and Ibrutinib, pyrrolidine versus piperidine (attached to 2-butynamide or acrylamide, respectively) as well as differences between the aromatic core structures of each drug. To begin to probe the molecular determinants of the observed differences in kinase domain dynamics and cellular efficacies, we tested the importance of the covalent bond between inhibitor and C481. Deuterium uptake changes in the wildtype BTK kinase domain C-lobe are abolished upon addition of either Tirabrutinib or Acalabrutinib to the BTK C481S mutant (Figure 2b). These data support the conclusion that the C481 mediated covalent bond is required to induce the allosteric effects observed in the C-lobe (FG loop/G-helix) of the BTK kinase domain prompting a closer examination of the different warheads.

### Swapping electrophilic warheads switches the dynamic allostery and functional profiles of BTK inhibitors

Two drug analogs, an Ibrutinib analog containing the Tirabrutinib warhead (2-butynamide-Ibrutinib) and a Tirabrutinib analog containing the Ibrutinib warhead (denoted acrylamide-Tirabrutinib), (Figure 3d, Supp. Fig. S3a)), were synthesized to determine the extent to which the different attachment modules affect potency in the B cell assay. The efficacy of each analog in terminating PLC_γ_2 phosphorylation and calcium signaling post BCR stimulation were measured as for the parent Ibrutinib and Tirabrutinib compounds. The 2-butynamide-Ibrutinib analog is less effective at inhibiting PLC_γ_2 phosphorylation as compared to Ibrutinib (Figure 3d,e). Additionally, the IC_50_ value for calcium flux inhibition of the 2-butynamide-Ibrutinib analog is similar to that of Tirabrutinib and the acrylamide-Tirabrutinib analog IC_50_ is similar to that of Ibrutinib (Figure 3f, Supp. Fig. 3a). To ensure that the differences in signaling efficacies were not due to different drug occupancies of BTK due to the differences in warhead reactivity, we measured BTK drug occupancy using a cellular thermal shift assay (CETSA) ^31^. As shown in Supp. Fig 3b,c, more soluble BTK is detected in the drug treated samples at higher temperatures compared to the DMSO treated control, suggesting that the drugs are binding and stabilizing BTK. Importantly, the relative levels of soluble BTK show no significant differences between Ibrutinib versus 2-butynamide-Ibrutinib analog treated cells, suggesting equivalent BTK drug occupancy in both samples. Taken together, swapping only the warheads between Tirabrutinib and Ibrutinib switches the efficacy of each drug.

We next reasoned that the observed switch in functional efficacy might correlate with the presence/absence of allosteric changes measured by HDX in the BTK kinase domain C-lobe (Figure 2a,b, Supp Fig. 1a,b). To test this idea, we compared the HDX profiles for Tirabrutinib, acrylamide-Tirabrutinib, Ibrutinib and 2-butynamide-Ibrutinib and find that only the 2-butynamide containing inhibitors cause increased deuterium uptake in the kinase domain C-lobe (Figure 4a). In contrast, all four inhibitors induce nearly identical HDX differences across the SH3-SH2 domain and surrounding the BTK active site (Figure 4a). As observed for the parent compounds, crystal structures of acrylamide-Tirabrutinib and 2-butynamide-Ibrutinib bound to the BTK kinase domain show no differences in the FG loop/G-helix region and covalent attachment is necessary; no changes in HDX are observed in the kinase domain C-lobe upon addition of acrylamide-Tirabrutinib or 2-butynamide-Ibrutinib to the BTK C481S mutant (Supp. Fig. S4a,b). Thus, the product of covalent attachment to the 2-butynamide warhead is necessary to induce allosteric changes in the BTK kinase domain C-lobe.

**Figure 4:**
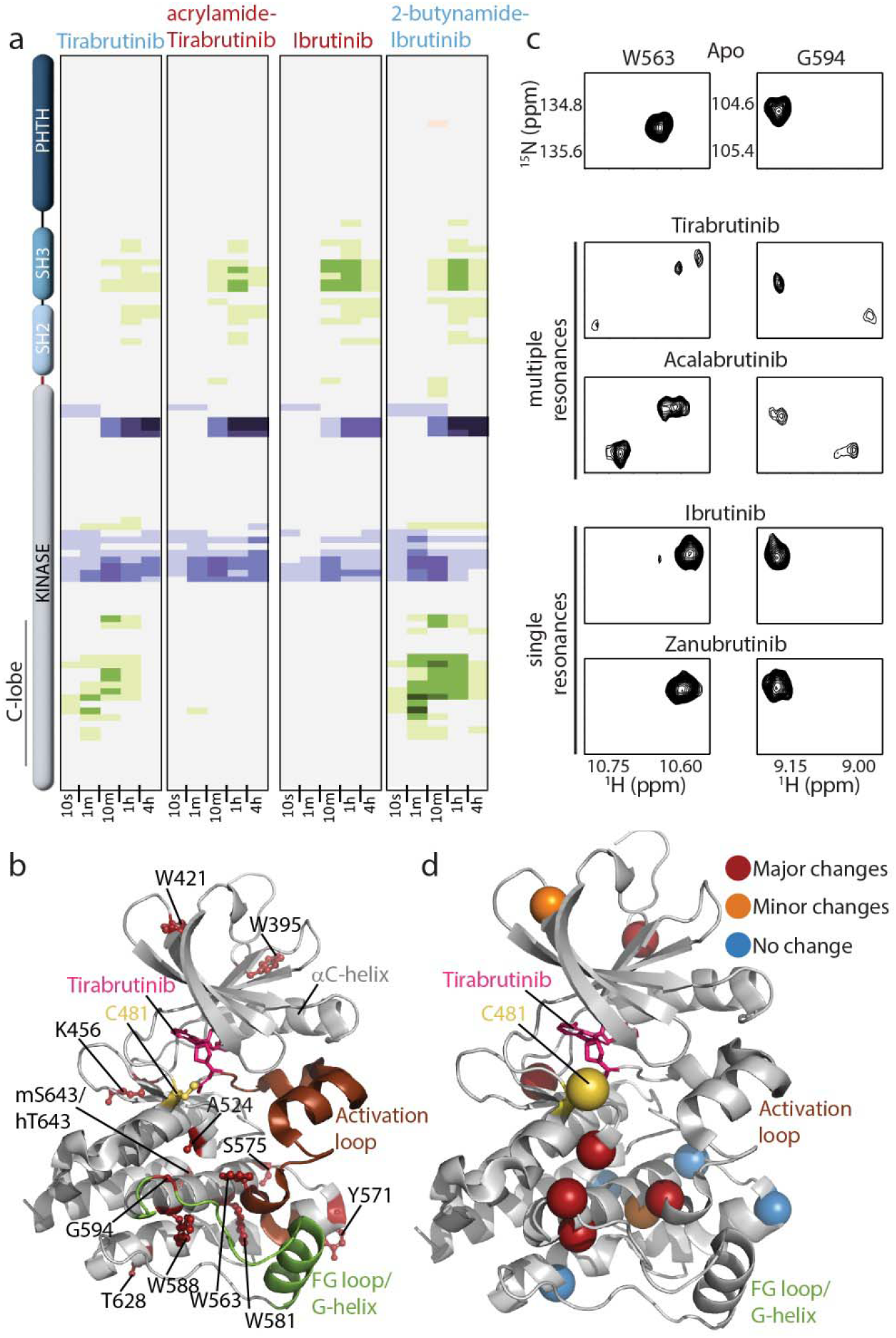
Swapping the electrophilic warhead between Tirabrutinib and Ibrutinib switches the solution behavior of the corresponding BTK/inhibitor complexes. (a) HDX-MS data for Tirabrutinib and Ibrutinib and corresponding analogs. (b) Residues with NMR assignments are shown as red sticks and labeled on the structure of Tirabrutinib bound BTK kinase domain. (c and d) Tirabrutinib or Acalabrutinib bound BTK adopt multiple conformations. (c) Select resonances from the ^1^H-^15^N TROSY-HSQC spectra of apo BTK kinase domain and four inhibitor complexes. (d) Kinase domain residues that give rise to multiple peaks upon binding of Tirabrutinib or Acalabrutinib are mapped onto the structure. Red spheres indicate well resolved multiple peaks observed for both drug/BTK complexes, orange are residues that show conformational heterogeneity in one or the other BTK/inhibitor, and blue represent residues that do not show multiple resonances.

To compliment the HDX-MS data, we applied NMR spectroscopic approaches to study the conformational effects of drug binding to the BTK kinase domain. Our previously reported backbone assignments for the apo BTK kinase domain ^32^ allowed unambiguous assignments for a subset of residues providing NMR probes distributed across the kinase domain (Figure 4b, Supp. Fig. S6). Clear differences in NMR spectral features emerge for the Tirabrutinib and Acalabrutinib complexes compared to the BTK kinase domain bound to Ibrutinib or Zanubrutinib. Binding of the 2-butynamide containing drugs (Tirabrutinib and Acalabrutinib) induces multiple NMR resonances for a subset of residues whereas the acrylamide containing inhibitors do not induce multiple resonances (Figure 4c, Supp. Fig. S6a). Conformational heterogeneity is abolished without the covalent bond between BTK and either Tirabrutinib or Acalabrutinib (Supp. Fig. S6b,c).

The appearance of multiple resonances for individual amino acids in the BTK kinase domain suggests slow conformational exchange and so we carried out ZZ-exchange experiments on BTK bound to Tirabrutinib. Off-diagonal peaks were detected providing direct evidence of exchange on the millisecond timescale (Supp. Fig. S6d). High-pressure NMR has also previously been used to characterize transient intermediate states ^33^. More broadly, hydrostatic pressure favors conformations with smaller partial molar volumes and can slow the rate of interconversion, thereby simplifying complex exchange processes ^34,35^. We therefore applied high-pressure NMR to the Tirabrutinib–BTK complex to define the multi-conformational states revealed by peak splitting and find that increasing pressure causes all of the split resonances to merge into single peaks at 2000–2500 bar, consistent with a dynamic equilibrium between multiple conformational states at ambient pressure (Supp Fig. S6e). The spectral changes observed at high pressure are reversible when the pressure is returned to 1 bar. By contrast, residues that do not exhibit peak splitting upon inhibitor binding exhibit only gradual chemical shift changes with pressure (Supp. Fig. S6e), an effect attributed to non-specific preferential hydration of the backbone that produces small downfield shifts universally observed in proteins under pressure ^36^. These observations confirm that slow conformational exchange is localized to specific residues of the Tirabrutinib-bound kinase domain, while the remainder of the domain behaves as a single, well-defined conformation.

Residues showing the most pronounced peak splitting are localized to the kinase domain C-lobe; the same region of the kinase domain that exhibit increased deuterium uptake by HDX-MS (FG loop/G-helix and base of activation loop) (Figure 4d). BTK residues in other regions of the kinase domain exhibit less pronounced spectral differences, either the presence of multiple peaks in spectra of only one and not both Tirabrutinib or Acalabrutinib bound BTK samples or appear as single peaks (Figure 4d, Supp. Fig S6a). Distinct conformational states of the FG loop/G-helix region could cause the observed increase in deuterium uptake in Tirabrutinib or Acalabrutinib bound BTK and in turn may drive the differences in substrate engagement and efficacy in terminating B cell signaling.

### Computational analysis of inhibitor binding to BTK

Having confirmed that 2-butynamide containing inhibitors induce multiple exchanging conformational states in the BTK kinase domain, we set out to use computational approaches to ascertain the source of the observed conformational heterogeneity. Analysis of over 100 x-ray structures of BTK bound to covalent active site inhibitors deposited in the Protein Data Bank shows that all but one exhibit the standard conformation associated with the inactive kinase activation loop (A-loop) (Figure 5a). One structure, that of Tirabrutinib bound to the BTK kinase domain, reveals a unique A-loop conformation, which we refer to as the ‘nonstandard’ inactive A-loop conformation (Figure 5b). The resolution of these two structures (1.08Å and 1.41Å, respectively) allows for a detailed description of inhibitor orientations. The tertiary amide of Ibrutinib (in the context of the standard inactive A-loop) adopts the cis isomer (Figure 5c,e). In contrast, the trans amide bond isomer is observed for Tirabrutinib whether the A-loop adopts the standard or non-standard conformation (Figure 5d,f and see PDB ID: 8FF0). In the latter, the trans amide isomer in Tirabrutinib permits formation of a hydrogen bond between the amide carbonyl of Tirabrutinib and the backbone amide NH of Q412 on the glycine-rich loop of the kinase domain (Figure 5d,f). This interaction allows hydrophobic packing of the adjacent BTK F413 sidechain into a pocket that is formed by the non-standard activation loop conformation (Figure 5d). The cis amide bond isomer observed in structures of Ibrutinib bound to BTK show no contacts between drug and the glycine-rich loop (Figure 5c,e).

**Figure 5:**
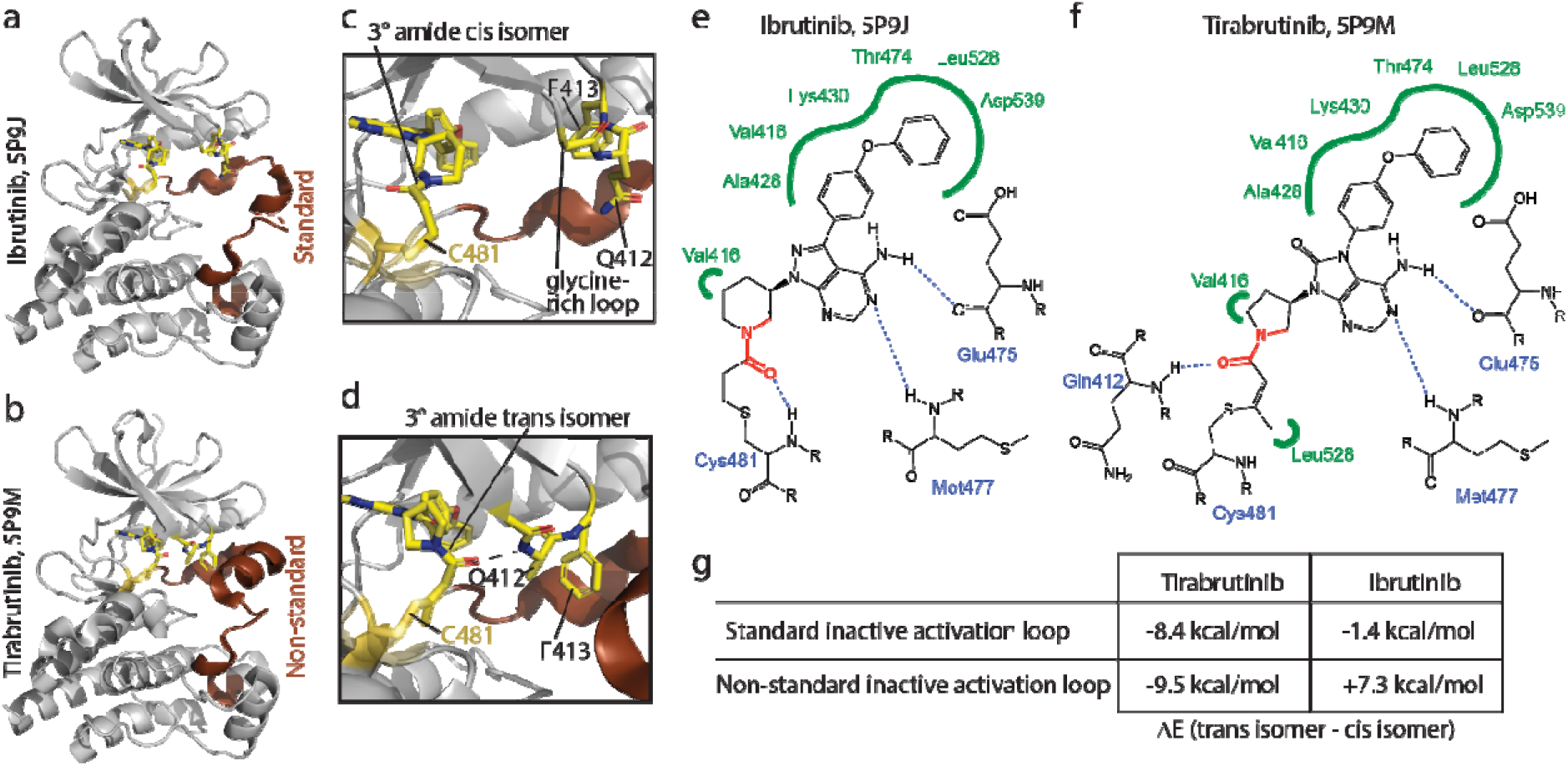
Density Functional Theory (DFT) calculations support the presence of multiple protein conformational states in Tirabrutinib bound BTK. (a and b) Structures showing the ‘standard’ and ‘non-standard’ inactive activation loop conformations in (a) Ibrutinib versus (b) Tirabrutinib bound BTK LKD. (c,d) Zoomed view of the BTK active site structure showing the cis amide bond isomer for Ibrutinib (c) and the trans amide bond isomer for Tirabrutinib (d). Also labeled are C481 and two residues (Q412 and F413) on the glycine rich loop. (e,f) 2D representations of (e) Ibrutinib and (f) Tirabrutinib binding poses adapted from Proteins*Plus*, PoseView and PoseEdit ^46^. The tertiary amide bond in each inhibitor is colored red. (g) Electronic energie obtained from DFT calculations are shown as ΔE values (kcal/mol) for Tirabrutinib and Ibrutinib in their cis versus trans amide bond configurations bound to the BTK kinase domain active sit with either the ‘standard’ or ‘non-standard’ inactive activation loop conformations. For each protein conformation, ΔE is defined as the calculated electronic energy of the inhibitor docked with the trans amide bond minus the energy of the inhibitor docked with the cis amide bond. The more thermodynamically favorable a given structure is, the more negative the energy value.

To further understand the interplay between inhibitor and protein conformational preferences, we generated quantum mechanics (QM)-based cluster models of the standard inactive A-loop and non-standard inactive A-loop consisting of ∼450 atoms surrounding the ATP binding site (Supp. Fig. S7). Tirabrutinib and Ibrutinib were docked to both enzyme conformations, in both the cis and trans amide-bond conformations (i.e. 4 states per inhibitor). Geometry optimization using Density Functional Theory (DFT) methods were performed and the energetics associated with each of the four states were compared (Figure 5g). For both A-loop conformations, the Tirabrutinib tertiary amide bond shows an energetic preference for the trans isomer in the BTK active site (Figure 5g). In contrast, the calculations suggest that the tertiary amide bond within Ibrutinib shows a marginal energetic preference for the trans isomer in the context of the standard inactive conformation of the A-loop and a clear preference for cis in the non-standard inactive A-loop conformation. Overall, these findings are consistent with the observation that Tirabrutinib and not Ibrutinib has been crystalized in both the standard and non-standard activation loop states.

From the computational experiments and available crystal structures a plausible structural mechanism explaining differences in C-lobe dynamics emerges wherein the tertiary amide bond of Tirabrutinib prefers to adopt the trans conformation within the ATP-binding site of BTK, irrespective of the A-loop conformation, while Ibrutinib can adopt both the cis and trans amide bond states. We postulate that the more limited conformational preference of the amide bond in Tirabrutinib may be the result of the adjacent alkene bond as opposed to the more flexible alkyl present in Ibrutinib/BTK (Figure 5e,f). Of course, like the tertiary amide bond, the alkene bond in the covalent adduct of Tirabrutinib (or Acalabrutinib) can also adopt Z and E isomers, albeit non-interconverting and determined by the reaction geometry. However, modeling the alkyne precursors of Tirabrutinib (or Acalabrutinib) into the BTK active site reveals only one orientation of the triple bond that allows for a productive nucleophilic attack by the C481 thiol. This is consistent with the observation that only the Z double bond isomer is captured in crystal structures of the covalent Tirabrutinib/Acalabrutinib BTK complexes and that the multiple conformations observed in solution are in dynamic exchange (Supp. Fig. S5d,e). Additionally, the methyl substituent on the resulting Z carbon-carbon double bond makes favorable hydrophobic contacts with the sidechain of L528 (Figure 5f); no such interactions are present in the Ibrutinib complex (Figure 5e). Collectively, we find that Tirabrutinib (and by extension Acalabrutinib) appear to adopt a single conformation in the active site of BTK (trans amide and Z alkene) and these 2-butynamide containing inhibitors stabilize at least two different BTK conformations (standard and non-standard A-loop) likely providing an explanation for the multiple NMR peaks observed for these complexes.

## DISCUSSION

The reliance of B cell cancers on BCR signaling has made BTK inhibition an effective treatment approach for specific B cell malignancies ^37,38^. The current collection of BTK inhibitors provide numerous treatment options resulting in complex clinical considerations in selection of specific treatment approaches. In addition, both Ibrutinib and Acalabrutinib are now approved for use in combination therapy with the BCL-2 inhibitor venetoclax with emerging discussions around which combination may be more effective ^39-41^. It will take time for long-term data to emerge on benefits/risks of one BTK inhibitor over another but it remains important to continue to advance our understanding of BTK inhibitor mechanism of action in the clinic, in B cells and at the molecular level.

The crystal structures of BTK in complex with the range of available covalent inhibitors are strikingly similar. It has therefore been assumed that the nature of the electrophile (and in turn the chemical structure of the covalent attachment to C481) does not affect the structure, dynamics or signaling capacity of inhibitor bound BTK. Our data demonstrate that covalent attachment of the 2-butynamide warhead on Tirabrutinib and Acalabrutinib to C481 (and not the acrylamide warhead) leads to allosteric changes in the dynamics and structure of the C-lobe of the BTK kinase domain. Tirabrutinib or Acalabrutinib bound BTK complexes both adopt multiple conformational states that are in slow exchange; this conformational heterogeneity correlates with an increase in binding to the BTK substrate PLCγ *in vitro*. The stronger association between drug bound BTK and PLCγ likely destabilizes the autoinhibited conformation of PLC_γ_ and increases the accessibility of PLC_γ_ for phosphorylation/activation by uninhibited BTK or other kinases ^21^. These differences are not sufficient to dramatically affect efficacy; all of these BTK inhibitors are effective in patients. It is however reasonable to imagine that the enhanced BTK/PLC_γ_ interaction and downstream signaling could create distinct opportunities for emergent mutations (in PLC_γ_ or other downstream molecules) that drive resistance mutations specific to the 2-buynamide warhead ^21,42^. Additional real-world data tracking resistance mutation profiles that occur under distinct drug pressures are needed to test this hypothesis.

The idea that differences in BTK inhibitor binding may drive differences in downstream signaling despite potent inhibition of BTK catalytic activity is borne out in the literature. In one study, Bender *et. al*., report that inhibitor binding mode (and the resulting conformation/phosphorylation status of the BTK kinase) results in differential compound potency against FcεR versus BCR signaling ^43^. A more recent study also focused on comparing inhibitors that occupy different regions of the active site (back pocket, hinge region and H3 pocket). In this work, Li *et al.* ^30^ examined the performance of different inhibitors in the context of acquired resistance. In an induced model of resistance, these authors found that Ibrutinib retained cytotoxic activity in the nanomolar range while Acalabrutinib (as well as the front pocket and hinge binders) did not ^30^. Li and co-workers also demonstrated that different BTK inhibitors trigger distinct downstream functional effects via non-catalytic scaffolding interactions. That work was predicated on our earlier biophysical studies showing that BTK inhibitors that bind the back pocket, hinge region or H3 (front) pocket cause distinct allosteric structural and dynamic effects in the remote non-catalytic regions of the BTK ^17,18^. Our results now show that even the closely related back pocket binders (Ibrutinib/Zanubrutinib versus Acalabrutinib/Tirabrutinib) trigger different conformational preferences in BTK due to differences in warhead structure.

Use of the acrylamide and 2-butynamide warheads is not limited to development of BTK inhibitors. Currently, at least eleven covalent inhibitors that use either the acrylamide or 2-butynamide warheads are approved for clinical use ^4,44^. These covalent inhibitors target a variety of proteins including G proteins such as KRAS and kinases such as epidermal growth factor receptor (EGFR), fibroblast growth factor receptor (FGFR) and Janus (JAK) kinases in addition to BTK ^44^. The success of these warheads has reinvigorated the search and development of other electrophiles for the design of covalent inhibitors. The focus of these development efforts has primarily been on optimizing reactivity and selectivity, as well as expanding chemoselectivity ^45^. Our findings suggest that different electrophilic warheads also impart distinct conformational/dynamic effects on their protein targets that can influence signaling outcomes.

## ACKNOWLEDGEMENTS

This work is supported by a grant from the National Institutes of Health (National Institute of Allergy and Infectious Diseases, AI43957) to A.H.A. and T.E.W., National Institutes of Health award 1S10OD032235 to A.H.A. to support the NMR infrastructure at Iowa State University and by funds from the Scott-Waudby Trust, the Hope Against Cancer charity, Cancer Research UK in conjunction with the UK Department of Health on an Experimental Cancer Medicine Centre grant [C10604/A25151] to R.G.B and M.J.SD. Research at the University of Leicester was carried out at the National Institute for Health and Care Research (NIHR) Leicester Biomedical Research Centre (BRC). The authors also thank the Roy J. Carver Charitable Trust, Muscatine, Iowa for ongoing research support. We thank Dr. Neha Amatya for her help in crystalizing the BTK kinase complexed with Ibrutinib. This research used resources of the NE-CAT beamline (GM124165) at the Advanced Photon Source (DE-AC02-06CH11357). The Eiger 16M detector on the 24-ID-E beam line is funded by a NIH-ORIP HEI grant (OD021527).

## AUTHOR CONTRIBUTIONS

R.E.J., R.G.B., M.J.S.D. and A.H.A. designed the experiments. R.E.J., R.G.B., T.E.W., D.Y.L., J.R., J.A.P., D.B.F., P.F., and M.P.G. performed the experiments. R.E.J., A.H.A., R.G.B., T.E.W., P.F., and M.P.G. analyzed the data. R.E.J., R.G.B., T.E.W., M.J.S.D., D.Y.L., J.R., J.A.P., D.B.F., P.F., M.P.G. and A.H.A. wrote and/or edited the manuscript.

## Abbreviations

BTK: Bruton’s Tyrosine Kinase
PRR: proline-rich region
LKD: linker-kinase domain
BCR: B cell receptor
PLC_γ_: Phospholipase C_γ_
HDX-MS: hydrogen/deuterium exchange-mass spectrometry
NMR: Nuclear Magnetic Resonance
WT: wild-type
DFT: Density Functional Theory
A-loop: Activation loop

**Supp. Data File 1**: Supplemental figures, table and methods. PDF file containing supplemental figures S1-S8, supplemental table 1, details of NMR chemical shift assignments and BTK inhibitor analog synthesis.

**Supp. Data File 2**: PDF file containing supporting NMR and MS data for acrylamide-Tirabrutinib, 2-butynamide-Ibrutinib and ‘alkane’ analog synthesis.

**Supp. Data File 3**: Excel file containing details of HDX-MS experimental set-up and differences in deuterium incorporation data for all peptides. Previously published HDX MS data have been included (taken from reference ^17^ identified with an asterisk) and those data have the dataset identifier PXD047865 in the PRIDE partner repository.

**Supp. Data File 4:** Excel file containing numerical data underlying the graphs used in the manuscript.

## REFERENCES

1 You, M., Liu, H. & Li, C. The Current Toolbox for Covalent Inhibitors: From Hit Identification to Drug Discovery. JACS Au 5, 5866–5887 (2025). 10.1021/jacsau.5c01134

2 Boike, L., Henning, N. J. & Nomura, D. K. Advances in covalent drug discovery. Nat Rev Drug Discov 21, 881–898 (2022). 10.1038/s41573-022-00542-z

3 Lu, X., Smaill, J. B., Patterson, A. V. & Ding, K. Discovery of Cysteine-targeting Covalent Protein Kinase Inhibitors. J Med Chem 65, 58–83 (2022). 10.1021/acs.jmedchem.1c01719

4 Zhao, Z. & Bourne, P. E. Exploring Extended Warheads toward Developing Cysteine-Targeted Covalent Kinase Inhibitors. J Chem Inf Model 64, 9517–9527 (2024). 10.1021/acs.jcim.4c00890

5 De Vita, E. 10 years into the resurgence of covalent drugs. Future Med Chem 13, 193–210 (2021). 10.4155/fmc-2020-0236

6 Rawlings, D. J. & Witte, O. N. The Btk subfamily of cytoplasmic tyrosine kinases: structure, regulation and function. Semin Immunol 7, 237–246 (1995). 10.1006/smim.1995.0028

7 Garg, N., Padron, E. J., Rammohan, K. W. & Goodman, C. F. Bruton’s Tyrosine Kinase Inhibitors: The Next Frontier of B-Cell-Targeted Therapies for Cancer, Autoimmune Disorders, and Multiple Sclerosis. J Clin Med 11 (2022). 10.3390/jcm11206139

8 Tavakoli, G. M., Yazdanpanah, N. & Rezaei, N. Targeting Bruton’s tyrosine kinase (BTK) as a signaling pathway in immune-mediated diseases: from molecular mechanisms to leading treatments. Adv Rheumatol 64, 61 (2024). 10.1186/s42358-024-00401-y

9 Shah, P. V. & Gladstone, D. E. Covalent and Non-Covalent BTK Inhibition in Chronic Lymphocytic Leukemia Treatment. Curr Treat Options Oncol 26, 754–763 (2025). 10.1007/s11864-025-01339-z

10 Lin, E. V., Arce, B., Alvarez-Arango, S. & Dispenza, M. C. Current and future landscape of Bruton tyrosine kinase inhibitors in allergy. J Allergy Clin Immunol 156, 568–578 (2025). 10.1016/j.jaci.2025.05.030

11. Klotz, L., Saraste, M., Airas, L. & Kuhlmann, T. Multiple sclerosis: 2024 update. Free Neuropathol 6, 14 (2025). 10.17879/freeneuropathology-2025-6762

12 Cui, Z. X. et al. Bruton’s tyrosine kinase (BTK) inhibitors: an updated patent review (2019-2024). Expert Opin Ther Pat, 1–25 (2025). 10.1080/13543776.2025.2554638

13 Kueffer, L. E., Joseph, R. E. & Andreotti, A. H. Reining in BTK: Interdomain Interactions and Their Importance in the Regulatory Control of BTK. Front Cell Dev Biol 9, 655489 (2021). 10.3389/fcell.2021.655489

14 Benrashid, S. & Woyach, J. A. When Targeted Therapy Falls Short: Unraveling Resistance Mechanisms in Chronic Lymphocytic Leukemia. Curr Hematol Malig Rep 21 (2026). 10.1007/s11899-026-00776-3

15 Rozkiewicz, D., Hermanowicz, J. M., Kwiatkowska, I., Krupa, A. & Pawlak, D. Bruton’s Tyrosine Kinase Inhibitors (BTKIs): Review of Preclinical Studies and Evaluation of Clinical Trials. Molecules 28 (2023). 10.3390/molecules28052400

16 Flanagan, M. E. et al. Chemical and computational methods for the characterization of covalent reactive groups for the prospective design of irreversible inhibitors. J Med Chem 57, 10072–10079 (2014). 10.1021/jm501412a

17 Joseph, R. E. et al. Impact of the clinically approved BTK inhibitors on the conformation of full-length BTK and analysis of the development of BTK resistance mutations in chronic lymphocytic leukemia. Elife 13 (2024). 10.7554/eLife.95488

18 Joseph, R. E. et al. Differential impact of BTK active site inhibitors on the conformational state of full-length BTK. Elife 9 (2020). 10.7554/eLife.60470

19 Joseph, R. E., Wales, T. E., Fulton, D. B., Engen, J. R. & Andreotti, A. H. Achieving a Graded Immune Response: BTK Adopts a Range of Active/Inactive Conformations Dictated by Multiple Interdomain Contacts. Structure 25, 1481–1494 e1484 (2017). 10.1016/j.str.2017.07.014

20 Joseph, R. E. et al. Activation loop dynamics determine the different catalytic efficiencies of B cell- and T cell-specific tec kinases. Sci Signal 6, ra76 (2013). 10.1126/scisignal.2004298

21 Joseph, R. E. et al. The Conformational State of the BTK Substrate PLCgamma Contributes to Ibrutinib Resistance. J Mol Biol 434, 167422 (2022). 10.1016/j.jmb.2021.167422

22. Quest Graph™ IC50 Calculator, AAT Bioquest, Inc., https://www.aatbio.com/tools/ic50-calculator (2026).

23 Johnson, B. A. & Blevins, R. A. NMR View: A computer program for the visualization and analysis of NMR data. J Biomol NMR 4, 603–614 (1994). 10.1007/BF00404272

24 Masson, G. R. et al. Recommendations for performing, interpreting and reporting hydrogen deuterium exchange mass spectrometry (HDX-MS) experiments. Nat Methods 16, 595–602 (2019). 10.1038/s41592-019-0459-y

25 Vizcaino, J. A. et al. 2016 update of the PRIDE database and its related tools. Nucleic Acids Res 44, 11033 (2016). 10.1093/nar/gkw880

26 Xie, Q., Joseph, R. E., Fulton, D. B. & Andreotti, A. H. Substrate recognition of PLCgamma1 via a specific docking surface on Itk. J Mol Biol 425, 683–696 (2013). 10.1016/j.jmb.2012.10.023

27 Min, L., Joseph, R. E., Fulton, D. B. & Andreotti, A. H. Itk tyrosine kinase substrate docking is mediated by a nonclassical SH2 domain surface of PLCgamma1. Proc Natl Acad Sci U S A 106, 21143–21148 (2009). 10.1073/pnas.0911309106

28 Li, W. et al. Bruton’s Tyrosine Kinase Inhibitors With Distinct Binding Modes Reveal Differential Functional Impact on B-Cell Receptor Signaling. Mol Cancer Ther (2023). 10.1158/1535-7163.MCT-22-0642

29 Montoya, S. et al. Kinase-impaired BTK mutations are susceptible to clinical-stage BTK and IKZF1/3 degrader NX-2127. Science 383, eadi5798 (2024). 10.1126/science.adi5798

30 Li, W. et al. Bruton’s Tyrosine Kinase Inhibitors with Distinct Binding Modes Reveal Differential Functional Impact on B-Cell Receptor Signaling. Mol Cancer Ther 23, 35–46 (2024). 10.1158/1535-7163.MCT-22-0642

31 Martinez Molina, D., et al. Monitoring drug target engagement in cells and tissues using the cellular thermal shift assay. Science 341, 84–87 (2013). 10.1126/science.1233606

32 Amatya, N. et al. Lipid-targeting pleckstrin homology domain turns its autoinhibitory face toward the TEC kinases. Proc Natl Acad Sci U S A 116, 21539–21544 (2019). 10.1073/pnas.1907566116

33 Zhang, Y. et al. High Pressure ZZ-Exchange NMR Reveals Key Features of Protein Folding Transition States. J Am Chem Soc 138, 15260–15266 (2016). 10.1021/jacs.6b09887

34 Roche, J. & Royer, C. A. Lessons from pressure denaturation of proteins. J R Soc Interface 15 (2018). 10.1098/rsif.2018.0244

35 Nguyen, T. T., Ghirlando, R., Roche, J. & Venditti, V. Structure elucidation of the elusive Enzyme I monomer reveals the molecular mechanisms linking oligomerization and enzymatic activity. Proc Natl Acad Sci U S A 118 (2021). 10.1073/pnas.2100298118

36 Williamson, M. P. Pressure-Dependent Conformation and Fluctuation in Folded Protein Molecules. Subcell Biochem 72, 109–127 (2015). 10.1007/978-94-017-9918-8_6

37 Honigberg, L. A. et al. The Bruton tyrosine kinase inhibitor PCI-32765 blocks B-cell activation and is efficacious in models of autoimmune disease and B-cell malignancy. Proc Natl Acad Sci U S A 107, 13075–13080 (2010). 10.1073/pnas.1004594107

38 Herman, S. E. et al. Bruton tyrosine kinase represents a promising therapeutic target for treatment of chronic lymphocytic leukemia and is effectively targeted by PCI-32765. Blood 117, 6287–6296 (2011). 10.1182/blood-2011-01-328484

39 Molica, S., Shanafelt, T. D., Giannarelli, D. & Allsup, D. Ibrutinib versus acalabrutinib in fixed-duration chronic lymphocytic leukemia therapy: a comparative analysis of efficacy. Haematologica 111, 412–417 (2026). 10.3324/haematol.2025.288323

40 Molica, S. & Allsup, D. Fixed-duration BTKi-venetoclax combinations in CLL: optimizing patient-centered care. Expert Rev Anticancer Ther 26, 295–299 (2026). 10.1080/14737140.2025.2590674

41 Mauro, F. R., Allsup, D. & Molica, S. Beyond mechanisms: toward patient-centered choices in BTKi-venetoclax fixed-duration therapy for CLL. Leukemia (2026). 10.1038/s41375-026-02945-y

42 Liu, T. M. et al. Hypermorphic mutation of phospholipase C, gamma2 acquired in ibrutinib-resistant CLL confers BTK independency upon B-cell receptor activation. Blood 126, 61–68 (2015). 10.1182/blood-2015-02-626846

43 Bender, A. T. et al. Ability of Bruton’s Tyrosine Kinase Inhibitors to Sequester Y551 and Prevent Phosphorylation Determines Potency for Inhibition of Fc Receptor but not B-Cell Receptor Signaling. Mol Pharmacol 91, 208–219 (2017). 10.1124/mol.116.107037

44 Roskoski, R., Jr. Orally effective FDA-approved protein kinase targeted covalent inhibitors (TCIs): A 2025 update. Pharmacol Res 217, 107805 (2025). 10.1016/j.phrs.2025.107805

45 Zhang, Z., Morstein, J., Ecker, A. K., Guiley, K. Z. & Shokat, K. M. Chemoselective Covalent Modification of K-Ras(G12R) with a Small Molecule Electrophile. J Am Chem Soc 144, 15916–15921 (2022). 10.1021/jacs.2c05377

46 Ehrt, C. et al. ProteinsPlus: a publicly available resource for protein structure mining. Nucleic Acids Res 53, W478–W484 (2025). 10.1093/nar/gkaf377

