## Supplemental Data File 1 for "BTK dynamics and function are differentially influenced by distinct covalent inhibitor warhead chemistries"

SUPPLEMENTAL FIGURES AND METHODS

SUPPLEMENTAL FIGURES:

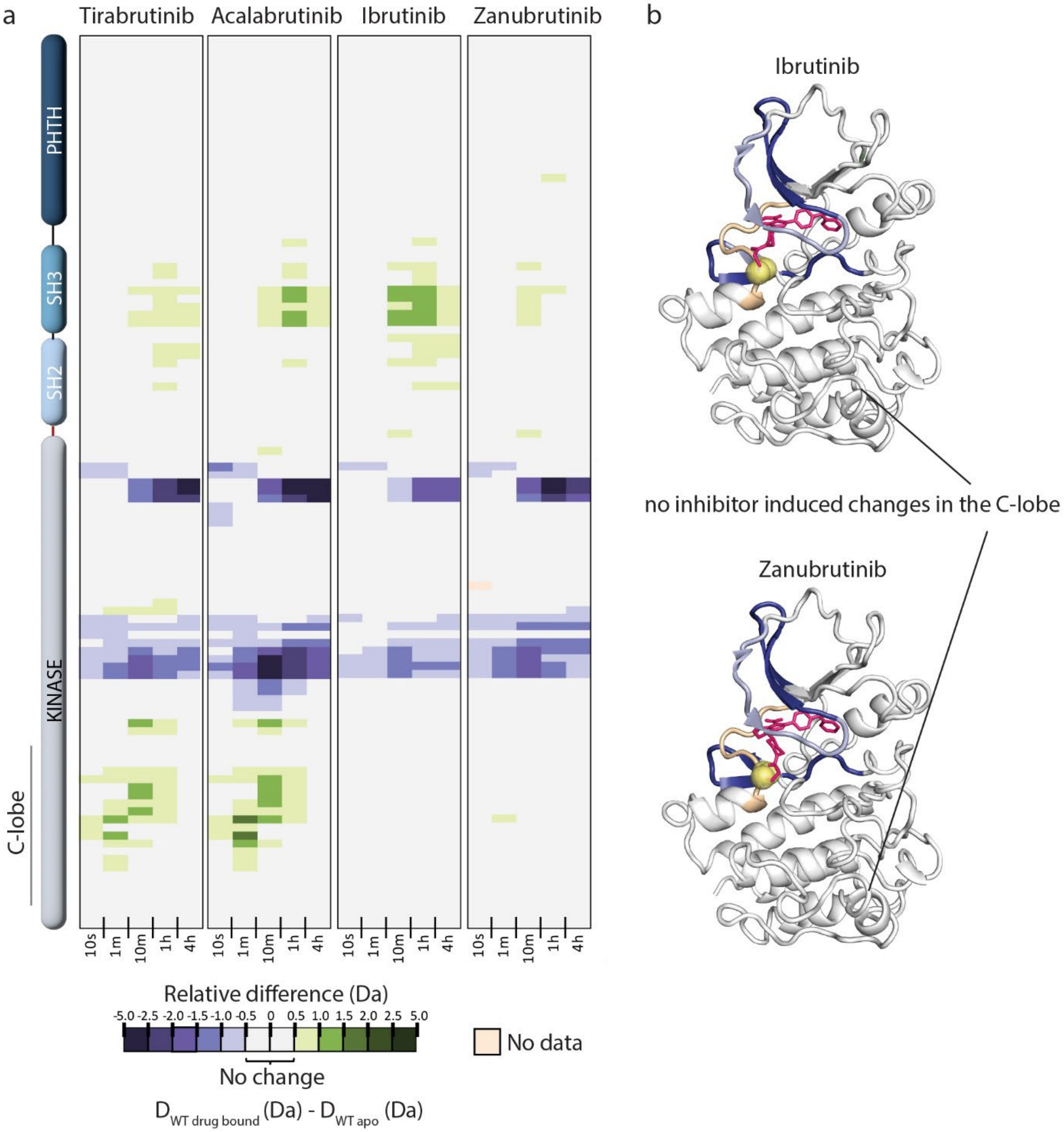


Supp. Fig. S1: (a) Tirabrutinib or Acalabrutinib bound BTK show increased deuterium uptake in the PLCγ substrate docking site (specifically FG loop/G-helix region of the kinase domain C-lobe) unlike Ibrutinib or Zanubrutinib bound BTK. Relative deuterium level of peptides in apo BTK was subtracted from the deuterium level of the corresponding peptide from each drug-bound form of BTK ($D_{WT\ drug\text{-}bound}$-$D_{WT\ apo}$) and the differences colored according to the scale shown. Peptic peptides are shown from N- to C-terminus, top to bottom, and the amount of time in deuterium is shown left to right. The relative difference data shown here represents a curated set of peptides that are coincident across all 4 states (apo and four drug-bound BTK forms). The identification of peptides, the relative difference values, and the complete data set for each state can be found in the Supp. Data File 3. The approximate position of the domains of BTK is shown at the left. (b) Changes in deuterium uptake in the BTK kinase domain upon binding to Ibrutinib and Zanabrutinib are show on the corresponding structures. Protection (shown in dark blue) occurs around the drug binding site. No changes are observed in the kinase domain C-lobe. Data for these inhibitors have been previously published and focused on the deuterium uptake changes in the N-terminal regulatory domains (1).

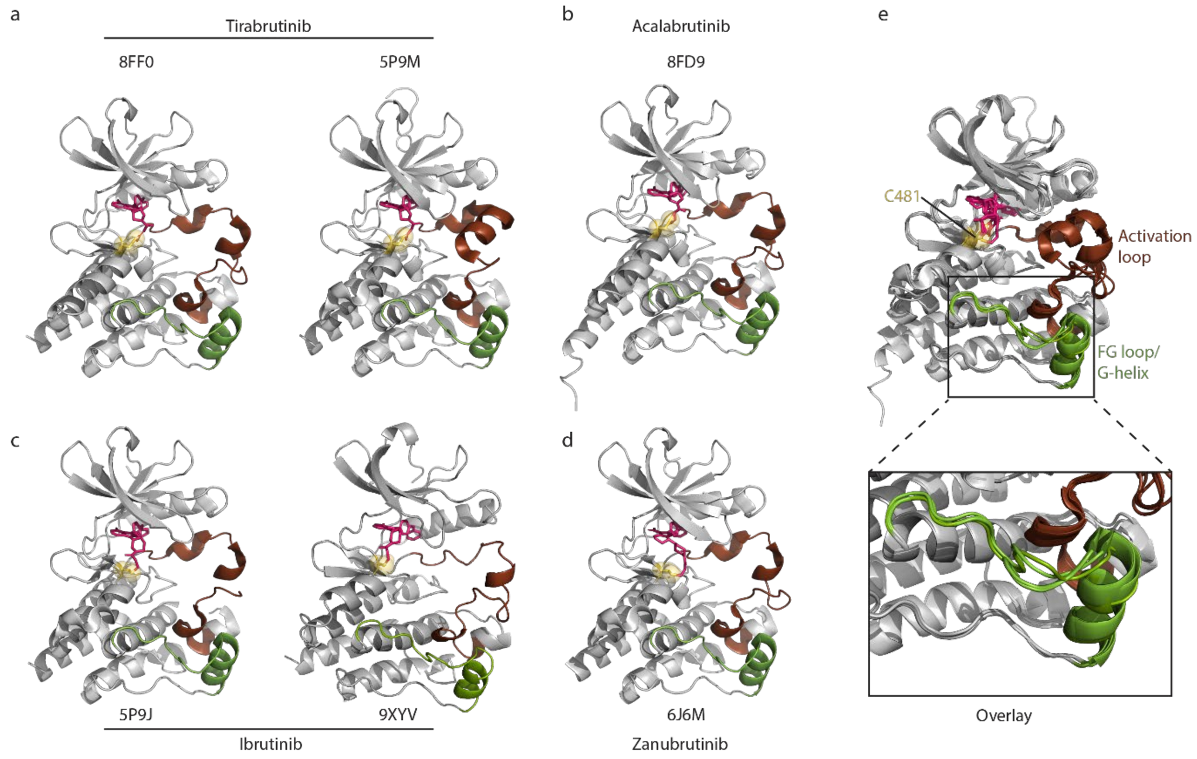


Supp. Fig. S2: Crystal structures of inhibitor bound BTK do not show significant differences in FG loop/G-helix. Crystal structures of BTK linker kinase domain bound to Tirabrutinib (PDB ID: 8FF0, 5P9M, (a)), Acalabrutinib (PDB ID: 8FD9), (b)), Ibrutinib (PDB ID: 5P9J, 9XYV, (c)) or Zanubrutinib (PDB ID: 6J6M), (d)). Colors are as in Figure 1. (e) An overlay of the structures in a-d shows no significant difference in the FG loop/G-helix (RMSD of 0.79 Å for the FG loop/G-helix (BTK L593-Q612) region). All structure figures were created using PyMOL (2).

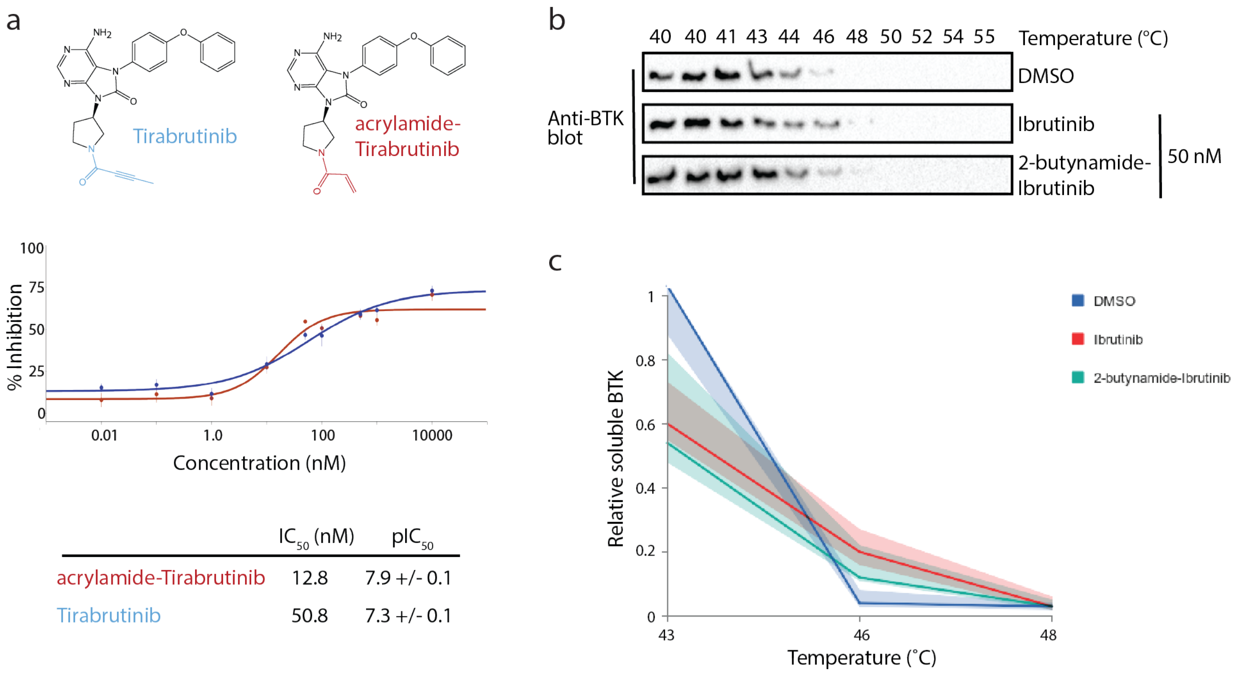


Supp. Fig. S3: The warhead on BTK inhibitors influences inhibitor effectiveness on downstream BCR signaling. (a) The acrylamide-Tirabrutinib analog more efficiently inhibits calcium flux post BCR stimulation of Ramos B cells compared to Tirabrutinib. Structures of both BTK inhibitors are shown highlighting the 2-butynamide (blue) and acrylamide (red) warheads. Half maximal inhibitory concentration ($IC_{50}$) values and $-log_{10}$ $IC_{50}$ ($pIC_{50}$) values ± SD (n=6) of calcium flux inhibition are listed. (b and c) Cellular thermal shift assay (CETSA) to test for BTK inhibitor occupancy. Western blot (b) and its quantification (details in methods) in (c) shows soluble BTK is detected at higher temperatures (46 and 48°C) in inhibitor treated samples compared to the DMSO control. Data from three independent experiments were averaged with the bold lines representing the median and the colored regions representing the interquartile range (IQR). Overlapping traces of Ibrutinib and 2-butynamide Ibrutinib suggests equivalent occupancy of BTK by either inhibitor. Source data for this figure is in Supp. Data File 4.

Methods:

*Cellular Thermal Shift Assay (CETSA)*: Ramos B cells (ATCC, # CRL-1596) were maintained in RPMI 1640 medium (Gibco, #A1049101) supplemented with 10% FBS (Thermo Fisher Scientific) and Penicillin/Streptomycin (Thermo Fisher Scientific) at 37°C/5% $CO_2$. Log phase growing cells were harvested, rinsed once with PBS and plated in RPMI 1640 medium without growth factors at a density of 1 x $10^6$ cells per well in a 12-well plate. The cells were incubated with 50 nM Ibrutinib, 2-butynamide-Ibrutinib or DMSO (0.1% final concentration) for one hour at 37°C/5% $CO_2$. The cells were harvested, rinsed twice in PBS with protease inhibitors at RT. The cell pellet was resuspended in 650 ul of PBS with protease inhibitors and placed in 50 ul aliquots in PCR tubes. The samples were heated for 3 min to varying temperatures (40 °C to 55 °C). The cells were cooled back to RT and then lysed by freeze-thawing. The lysed cells were spun at 14K for 12 min at 4 °C. The supernatants were boiled with SDS-PAGE dye and analyzed by Western immunoblot analysis using an Anti-BTK antibody (BD Transduction Laboratories, # 611117). The intensity of the bands was quantified using Image Lab (Biorad). The intensity of the BTK band at 40 °C was set to one and the intensity of BTK bands at the higher temperatures were reported relative to this. Data from three independent experiments were averaged and reported as median with interquartile range (IQR).

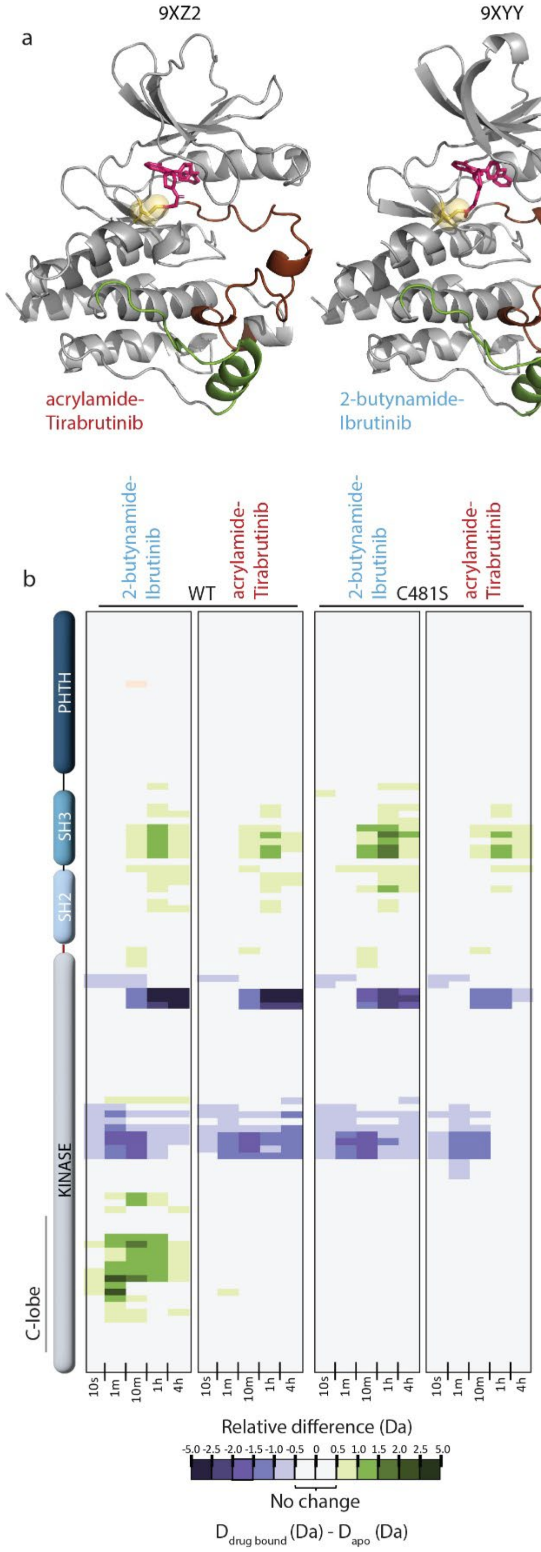


Supp. Fig. S4: (a) Crystal structures of acrylamide-Tirabrutinib (left, PDB ID: 9XZ2) and 2-butynamide-Ibrutinib (right, PDB ID: 9XYY) bound to the BTK kinase domain. No significant differences in the FG loop /G-helix were observed when compared to their parent compound-BTK LKD co-complexes (RMSD of 0.82 Å and 1.12 Å respectively). (b) Covalent attachment between the 2-butynamide warhead and BTK C481 is required for the increased deuterium exchange observed in the FG loop/G-helix (C-lobe region) of BTK. Mutation of C481 to serine abolishes allosteric effects in the kinase domain C-lobe.

Methods:

*Crystallization, structure determination, and refinement:*

Purified BTK KD K430R-Loop swap mutant protein at 18 mg/ml in 20 mM Tris-HCl, (pH 8), 150 mM NaCl, and 10% glycerol were mixed with 1 mM of the respective inhibitor and 10% DMSO on ice for 30 minutes. Crystals of the complexes were obtained by mixing protein/inhibitor solutions with precipitant solutions at a 1:1 ratio using the sitting-drop diffusion method at 4 °C. The BTK KD/Ibrutinib complex was crystallized in 20% PEG 3350 and 0.1M sodium citrate, (pH 5.5); the BTK KD/2-butynamide-Ibrutinib complex was crystallized in 25% PEG 3350, 0.1M Bis-Tris, (pH 5.5), and 0.2M ammonium sulfate; the BTK KD/acrylamide-Tirabrutinib complex was crystallized in 20% PEG 4000 and 0.1M sodium citrate, (pH 5.6). Crystals were harvested in cryoprotectant solutions consisting of the respective crystallization conditions supplemented with 20% glycerol and flash-frozen in liquid nitrogen.

X-ray diffraction data were collected at APS beamline 24-ID-E (NE-CAT). Data sets were indexed, integrated, and scaled using *autoPROC* (Global Phasing) (3-5). The structures of protein/inhibitor complexes were solved by molecular replacement using *PHASER*, (6). Three-dimensional structures of the inhibitors were generated using Grade Web Server, (7) and the ligand restraints were prepared with *eLBOW* (8). Model building was performed using Coot (9), and structure refinement was carried out with Phenix (10). The twin law of –h, –k, l was used in the refinement of BTK KD/acrylamide-Tirabrutinib structure due to observed crystal twinning. Crystallographic data collection and refinement statistics are listed in Supp. Table 1. The atomic coordinates and structure factors have been deposited into the PDB with the accession codes: 9XYV (BTK KD/Ibrutinib), 9XYY (BTK KD/2-butynamide-Ibrutinib), and 9XZ2 (BTK KD/acrylamide-Tirabrutinib).

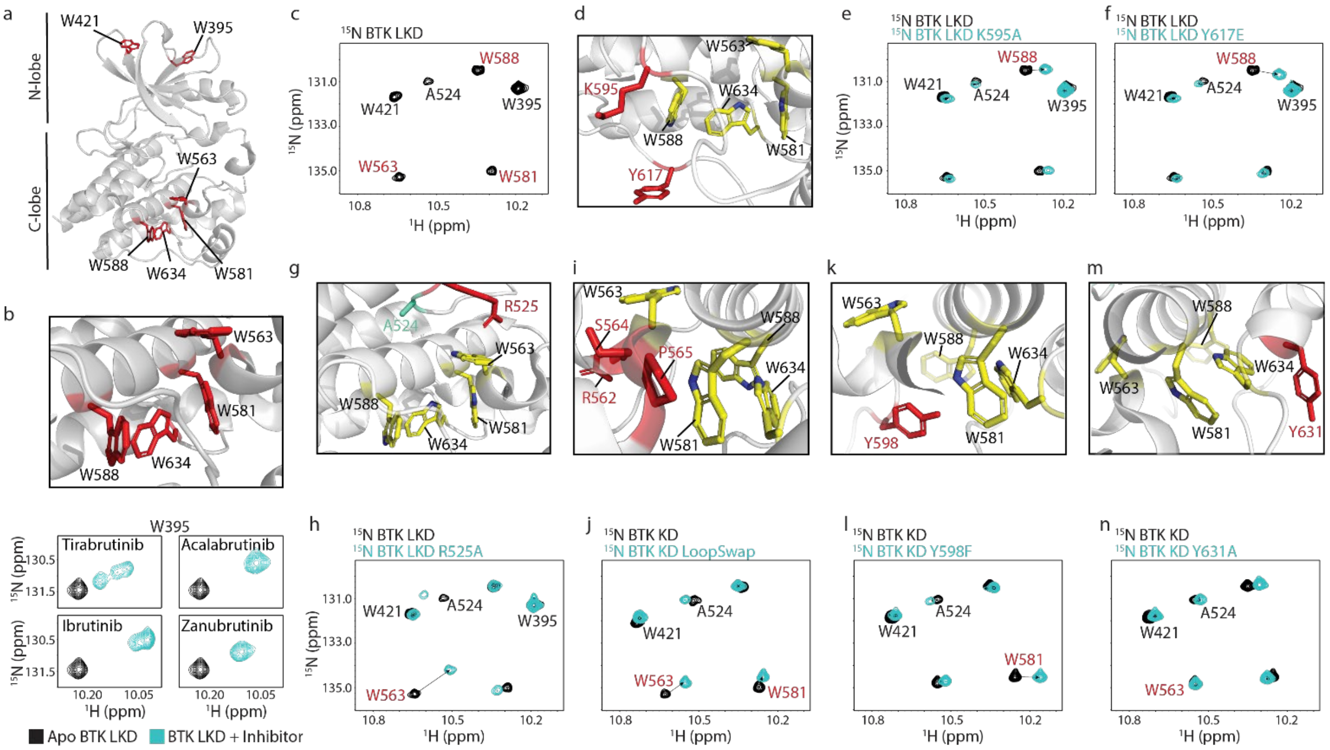


Supp. Fig. S5: Assignment of tryptophan residues in the C-lobe of BTK kinase. (a) Structure of BTK linker kinase domain (LKD) with all the tryptophan residues incidated as red sticks. The kinase N-lobe tryptophan residues (BTK W395 and W421) have been previously assigned (11). (b) Close up of the kinase C-lobe tryptophan residues. Bottom: Spectral overlay of the $^{1}$H-$^{15}$N TROSY HSQC spectra of apo and inhibitor bound kinase domain for each of the four clinically relevant BTK inhibitors show chemical shift changes consistent with inhibitor binding to BTK and indicate that the kinase domain is fully occupied by inhibitor; no resonances are detectable for the unbound apo kinase. (c) The tryptophan indole side chain region of the $^{1}$H-$^{15}$N TROSY-HSQC spectrum of $^{15}$N-labelled BTK LKD. Of the six tryptophan residues in BTK LKD, only 5 are visible in the $^{1}$H-$^{15}$N TROSY-HSQC spectrum. Previously assigned peaks are labeled in black. Peaks assigned in this work are labeled in red. (d, g, i, k and m) Structures of BTK showing residues that are mutated as red sticks. Tryptophan residues to be assigned are shown as yellow sticks. (e, f, h, j, l and n) $^{1}$H-$^{15}$N TROSY-HSQC spectral overlay of the indole NH region of BTK LKD or BTK kinase domain (KD) WT (black) with BTK mutants (cyan). Peaks showing major chemical shift changes are indicated with an arrow and labeled in red.

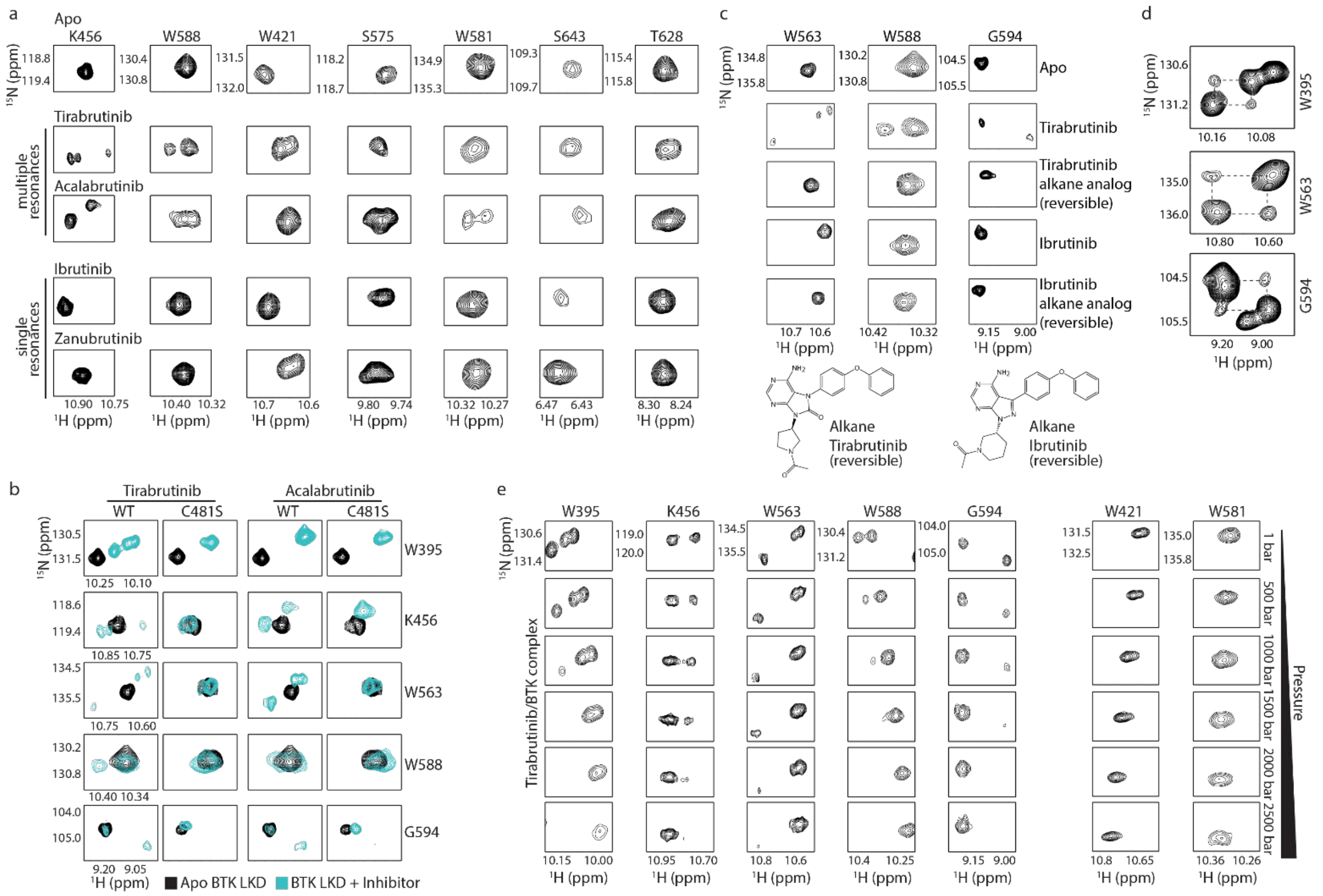

Supp. Fig. S6: The different inhibitor bound BTK complexes give rise to distinct NMR spectra. (a) Spectral changes induced by binding of Tirabrutinib, Acalabrutinib, Ibrutinib or Zanubrutinib to $^{15}N$-labeled BTK LKD are shown for select resonances. Examples of peak multiplicity in the Tirabrutinib and Acalabrutinib complexes (but not Ibrutinib and Zanubrutinib complexes) are shown in addition to residues for which only a single resonance is observed for any drug/BTK complex (S643, T628). (b) $^{1}H$-$^{15}N$ TROSY-HSQC spectral overlay of select BTK resonances in the apo (black) and inhibitor bound (cyan) form. Multiple peaks that are observed upon inhibitor binding for resonances in the BTK WT spectrum are absent in the BTK C481S mutant. (c) Complimenting the C481S mutation in panel (b), use of a non-reactive (reversible) alkane analogs of Tirabrutinib and Ibrutinib show no evidence of multiple species by NMR upon binding the BTK kinase domain. Data are as described in (a). (d) Off-diagonal cross-peaks observed for BTK W395, W563 and G594 resonances in ZZ exchange experiment conducted on Tirabrutinib bound BTK LKD sample. (e) $^{1}H$-$^{15}N$ TROSY-HSQC spectra for residues in the Tirabrutinib/BTK complex at increasing pressures (1 bar to 2500 bar). High pressure causes multiple resonances to merge into a single peak.

Methods:

*High pressure NMR*: 350 µl of uniformly $^{15}N$ labeled BTK LKD (300 µM) in NMR buffer containing 400 µM Tirabrutinib and 2% DMSO was loaded into a commercially available high-pressure 5 mm zirconia NMR tube. After sample loading the tube was topped with mineral oil to form an immiscible layer, allowing the application of external pressure. The filled NMR tube was connected to a Daedalus Xtreme-60 high pressure automated syringe pump (Daedalus Innovations, Philadelphia, PA) which varied the pressure from 1-2500 bar. Multiple $^{1}H$-$^{15}N$ TROSY-HSQC spectra at varying pressure were collected in 500 bar increments at 298 K.

*ZZ exchange NMR*: The $^{1}H$-$^{15}N$ ZZ-exchange NMR experiment was carried out on a 300 µM uniformly $^{15}N$ labeled BTK LKD with 400 µM Tirabrutinib in NMR buffer with 2% DMSO at 500 bar and 298 K. The experiment was acquired with 160*($t_1$, $^{15}N$) x 2048*($t_2$, $^{1}H$) complex points, a 1.5 interscan delay (D1), and a 600 ms $t_m$ (mixing time). The sample was maintained at 500 bar (instead of ambient pressure) for enhanced sample stability at higher pressure.

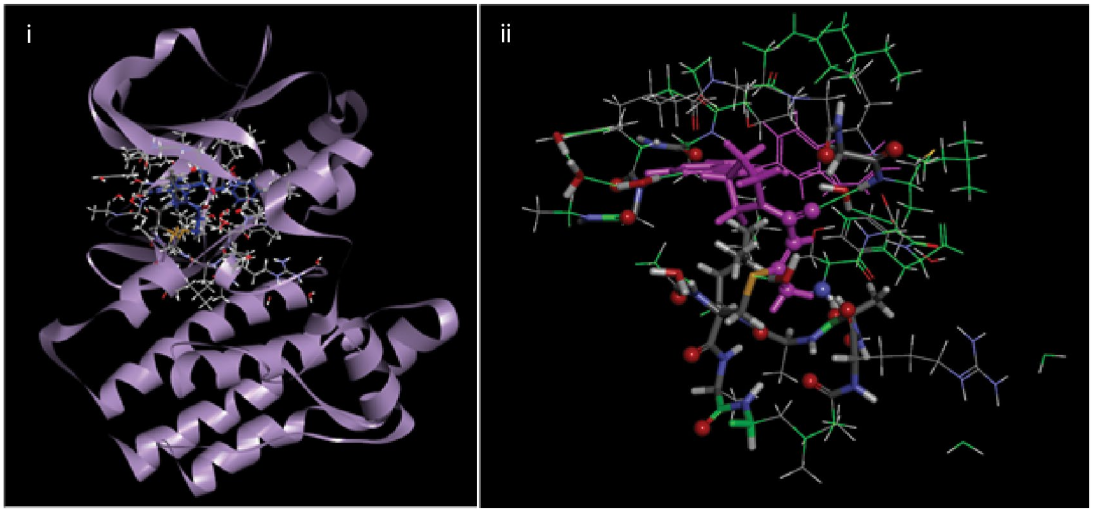


Supp. Fig. S7: Illustration of the QM models used in this study. (i) model shown in relation to the kinase domain and (ii) atoms included in the model. Pink colors indicate the bound inhibitor, green indicates fixed protein atoms, sticks molecular representations indicate atoms optimized at M06-2X/6-31G(d) and lines indicate atoms optimized at M06-2X/6-31G.

Methods:

*Density function theory calculations*:

A QM model representing the local ATP binding site found within 5 Å of the BTK inhibitors was created from the 5P9M and 5P9J structures. The QM subsystem consisted of 390 atoms representing the enzyme and 8 conserved water molecules. An additional ~60 atoms were needed to model Tirabrutinib and Ibrutinib. The QM region includes the side chains or backbone atoms from residues; Leu408-Gln412, Val416, Ala428, Lys430, Met449, Val458, Ile472, Thr474-Ala478, Gly480-Leu483, Ala524-Cys527, Leu528, Ser538-Phe540 and Leu542. To maintain the shape of the pocket 60 of the ~450 atoms were frozen (Supp. Fig. S7). All geometry optimization calculations were performed in the Gaussian 16 program (12). Structures were optimized using the

M06-2X functional of Truhlar and co-workers (13-15) in a polarized continuum solvent model consisting of water. Approximately 190 of the atoms, including all those involved in direct enzyme-inhibitor H-bonding, were treated using the 6-31G(d) basis set. Approximately 280 more distant atoms, or sidechains interacting via VDW interactions, were treated using the 6-31G basis set (Supp. Fig. S7). All optimized stationary points were confirmed based on their vibrational frequencies. We report the difference in the electronic energies for the same inhibitor in two different enzyme conformations to assess whether either Ibrutinib or Tirabrutinib have a particular preference for either form.

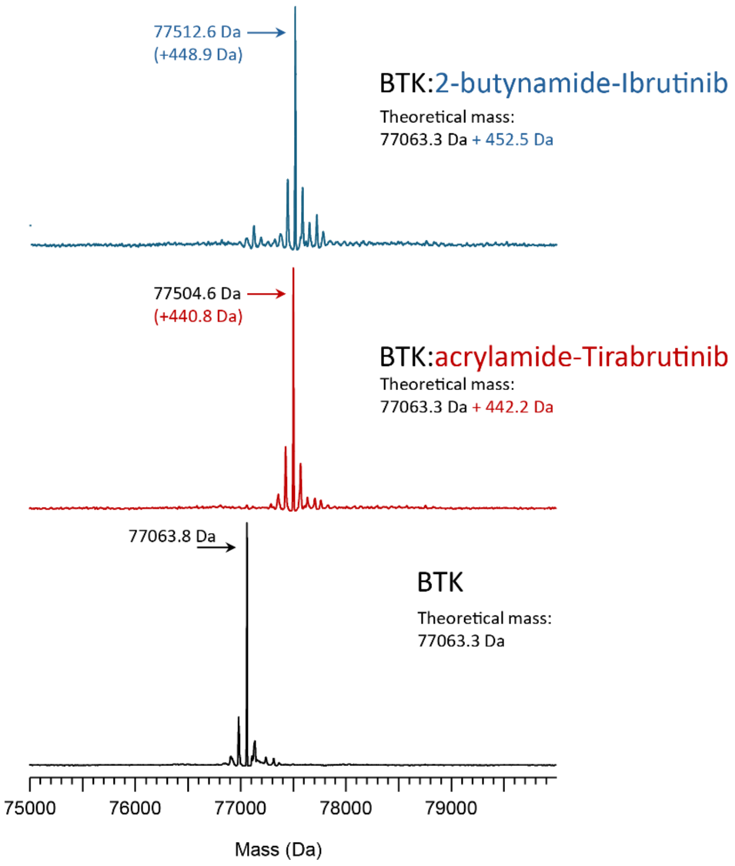


Supp. Fig. S8: Intact mass analysis of apo and inhibitor bound BTK full-length (FL). BTK FL was incubated for one hour with a twofold excess of 2-butynamide-Ibrutinib (blue spectrum) or acrylamide-Tirabrutinib (red spectrum). Intact mass analysis shows a mass increase of one covalent inhibitor molecule per BTK molecule compared to apo BTK FL (black spectrum).

Supplemental Table 1: Crystallographic data collection and refinement statistics.

| Supplemental Table 1: Crystallographic data collection and refinement statistic | | | |
|---|---|---|---|
| | **BTK KD in complex with Ibrutinib** | **BTK KD in complex with 2-butynamide-Ibrutinib analog** | **BTK KD in complex with acrylamide-Tirabrutinib analog** |
| **PDB code** | 9XYV | 9XYY | 9XZ2 |
| **Data collection** | | | |
| Resolution range§ | 93.02 - 1.34 (1.338 - 1.342) | 93.18 - 2.43 (2.79 - 2.43) | 53.28 - 2.48 (2.50 - 2.48) |
| Space group | P 61 | P 61 | P 31 |
| Cell dimensions | | | |
| a, b, c (Å) | 107.41 107.41 45.15 | 107.59 107.59 45.96 | 106.55 106.55 46.38 |
| *α, β, γ (°)* | 90 90 120 | 90 90 120 | 90 90 120 |
| Multiplicity§ | 9.9 (4.0) | 8.1 (7.4) | 16.8 (4.0) |
| Completeness (%)§ | 99.49 (89.74) | 90.0 (35.4) | 100.0 (100.0) |
| Mean I/sigma (I) § | 8.30 (0.74) | 9.7 (1.2) | 13.61 (1.45) |
| R-merge§ | 0.14 (1.44) | 0.131 (2.402) | 0.150 (0.154) |
| $CC_{1/2}$§ | 0.995 (0.333) | 0.928 (0.317) | 0.998 (0.379) |
| **Refinement** | | | |
| Resolution range§ | 40.61 - 1.37 (1.39 - 1.37) | 53.80 – 2.48 (2.57 - 2.48) | 53.28 - 2.48 (2.59 - 2.48) |
| Unique reflections§ | 60884 (2935) | 9543 (638) | 20664 (2372) |
| R-work§ | 0.1424 (0.2348) | 0.2106 (0.3367) | 0.2334 (0.3342) |
| R-free§ | 0.1689 (0.3347) | 0.2414 (0.39.23) | 0.2814 (0.3621) |
| Non-hydrogen atoms (n) | 2550 | 2103 | 4069 |
| Protein | 2182 | 2052 | 4003 |
| Ligand | 59 | 34 | 66 |
| Water | 309 | 17 | 0 |
| Protein residues | 265 | 262 | 265 |
| Average B-factors ($Å^2$) | 19.18 | 74.45 | 88.07 |
| Protein | 17.40 | 74.79 | 88.31 |
| Ligand | 26.01 | 62.51 | 73.18 |
| Water | 30.41 | 57.00 | - |
| RMS bonds (Å) | 0.003 | 0.004 | 0.002 |
| RMS angles (°) | 0.74 | 0.87 | 0.60 |
| Ramachandran favored (%) | 98.86 | 98.84 | 98.48 |
| Ramachandran allowed (%) | 1.14 | 1.16 | 1.52 |
| Ramachandran outliers (%) | 0.00 | 0.00 | 0.00 |

§ Statistics for the highest resolution shell are shown in parentheses.

*Assignment of tryptophan side chains NMR resonances in BTK LKD:*

The BTK LKD contains six tryptophan residues: two in the N-lobe of the kinase domain, (W395 and W421) and four in the C-lobe of the kinase (W563, W581, W588 and W634), (Supp. Fig. S5a,b). The indole NH side chain region of the $^{1}$H-$^{15}$N TROSY-HSQC spectrum of BTK LKD WT has six cross-peaks (Note: BTK 'WT' contains the previously described BTK Y617P mutation required for the expression of soluble BTK protein in bacteria, (11)) (Supp. Fig. S5c). Of the six cross-peaks in the $^{1}$H-$^{15}$N TROSY-HSQC spectrum of BTK LKD WT, we have previously assigned three cross-peaks: two cross-peaks correspond to the kinase N-lobe tryptophan residues (W395 and W421) and the third cross-peak corresponds to a non-tryptophan residue: A524 (11, 16). This leaves four unassigned tryptophan residues in the C-lobe of the kinase (W563, W581, W588 and W634) and three cross-peaks. This suggests that one of the remaining four C-lobe tryptophan residues is likely broadened beyond detection in the BTK LKD WT spectrum.

To assign the C-lobe tryptophan residues of BTK kinase, we individually mutated each tryptophan residue individually to alanine. However, we were unable to express soluble protein for any of the alanine mutants. We additionally tried mutating each of the tryptophan residues to phenylalanine or valine, but none of these mutants were expressed as soluble protein. Next, we tried a proximity approach to assign the C-lobe tryptophan residues. Mutation of a residue adjacent to a given tryptophan is expected to cause large chemical shift changes in the resonance corresponding to that tryptophan with little or no changes to residues that are further away.

BTK K595 and Y617 are adjacent to W588 and further away from W563, W581 and W634 (Supp. Fig. S5d). Mutation of BTK K595 and Y617 are expected to cause major chemical shift changes in W588. As shown in the spectral overlay of the tryptophan side chain region of BTK LKD WT spectrum (black spectrum) with that of the BTK LKD K595A or Y617E mutant spectrum (in cyan), mutation of either BTK K595 or Y617 induces a major chemical shift change in one cross-peak and little or no change in the other cross-peaks (Supp. Fig. S5e,f). The cross-peak that showed the major chemical shift change was tentatively assigned as W588.

BTK R525 is in close proximity to W563 and further away from the other three C-lobe tryptophan residues (Supp. Fig. S5g). As shown in Supp. Fig. S5h, a major chemical shift change is observed in one of the unassigned tryptophan cross-peaks and a minor change or no change in the other unassigned cross-peaks. Additionally, as expected, BTK A524 which is adjacent to R525 undergoes a major chemical shift change. The unassigned cross-peak that underwent the major chemical shift change was therefore tentatively assigned as W563.

BTK R562, S564 and P565 in the BTK kinase activation loop are in close proximity to BTK W563 and W581(Supp. Fig. S5i). We have previously mutated BTK R562, S564 and P565 residues along with three other loop residues (L542, S543 and V555) to the corresponding loop residues in ITK (referred to as the LoopSwap mutations: L542M, S543T, V555T, R562K, S564A and P565S, (17)). Comparison of the $^{15}$N TROSY-HSQC spectra of BTK LKD with and without the LoopSwap mutations (Supp. Fig. S5j), shows major chemical shift changes in two cross-peaks: one was the putative W563 cross-peak and the other cross-peak that we now tentatively assign as W581.

To further confirm the assignment of W581, we mutated BTK Y598 which is adjacent to W581 and further away from the other tryptophan residues (Supp. Fig. S5k). An overlay of the $^{15}$N TROSY-HSQC spectra of BTK LKD with and without the Y598F mutation shows a major chemical shift change in the cross-peak that we had tentatively assigned as W581 (Supp. Fig. S5l).

BTK W581 and W588 are located on a stable secondary structure within the kinase: the F-helix, whereas, BTK W563 and W634 are located in potentially more dynamic regions within the kinase: the base of the kinase activation loop (W563) or a loop that interconnects helices H and I (W634) (Supp. Fig. S5b, top). Hence, BTK W563 or W634 are likely candidates to be broadened beyond detection in the $^{15}$N TROSY-HSQC spectrum of BTK LKD WT. BTK W563, W581 and W588 have been tentatively assigned to the three cross-peaks in the $^{15}$N TROSY-HSQC spectra of BTK LKD WT. By elimination, BTK W634 is therefore expected to be the resonance that is broadened beyond detection in the $^{15}$N TROSY-HSQC spectrum of BTK LKD WT. To confirm these assignments, BTK Y631 which is close to W634 was mutated to alanine (Supp. Fig. S5m). As shown in the overlay of the BTK WT and Y631A mutant spectra (Supp. Fig. S5n), only minor chemical shift changes are observed in all the cross-peaks, suggesting that W634 is likely not visible in the $^{15}$N TROSY-HSQC spectrum of BTK LKD WT. Taken together, BTK W395, W421, W563, W581 and W588 indole NH are visible and assigned in the $^{15}$N TROSY-HSQC spectrum of BTK LKD WT.

*Analog synthesis*:

**(*R*)-3-(4-phenoxyphenyl)-1-(piperidin-3-yl)-1*H*-pyrazolo[3,4-d]pyrimidin-4-amine.HCl**

Boc-protected Ibrutinib intermediate (*tert*-butyl (*R*)-3-(4-amino-3-(4-phenoxyphenyl)-1*H*-pyrazolo[3,4-*d*]pyrimidin-1-yl)piperidine-1-carboxylate (BLDpharma, BD560719), 700 mg, 1.44 mmoles) was dissolved in MeOH/ EtOAc (1:1, 40 mL), then a solution of HCl in dioxane (4M, 4.0 mL, 16 mmoles) was added and stirred at RT for 24 hrs. Volatiles were removed in vacuo, the resulting white solid triturated with hexane (3 x 5 mL) and dried to yield product as a white solid (605 mg). 99% Yield (Supp. Data File 2).

**(*R*)-1-(3-(4-amino-3-(4-phenoxyphenyl)-1*H*-pyrazolo[3,4-*d*]pyrimidin-1-yl)piperidin-1-yl)but-2-yn-1-one**

But-2-ynoyl chloride was prepared in situ: 2-Butynoic acid (50 mg, 0.59 mmoles) was dissolved in DCM (5 mL) and cooled to 0 °C. A solution of oxalyl chloride in DCM (2M, 295 µL, 0.59 mmoles) was added followed by DMF (~50 µL), then stirred at RT for 2 hrs.

(*R*)-3-(4-phenoxyphenyl)-1-(piperidin-3-yl)-1*H*-pyrazolo[3,4-*d*]pyrimidin-4-amine.HCl (250 mg, 0.59 mmoles) was suspended in DCM (20 mL) and cooled to 0 °C. DIPEA (310 µL, 229 mg, 1.78 mmoles) was added followed by but-2-ynoyl chloride (prepared above). The reaction mixture was stirred at 0 °C for 30 mins, then MeOH (1 mL) was added dropwise and stirred for 10 mins before concentrating in vacuo. Flash chromatography purification as performed (silica, 2% → 5%MeOH in DCM) to yield a white solid (227 mg). 85% yield. (Supp. Data File 2).

**(*R*)-1-(3-(4-amino-3-(4-phenoxyphenyl)-1*H*-pyrazolo[3,4-*d*]pyrimidin-1-yl)piperidin-1-yl)ethan-1-one**

(*R*)-3-(4-phenoxyphenyl)-1-(piperidin-3-yl)-1*H*-pyrazolo[3,4-*d*]pyrimidin-4-amine.HCl (250 mg, 0.59 mmoles) was suspended in DCM (20 mL) and cooled to 0 °C. DIPEA (310 µL, 229 mg, 1.78 mmoles) was added followed by acetyl chloride (42 µL, 46 mg, 0.59 mmoles). The reaction mixture was stirred at 0 °C for 30 mins, then MeOH (1 mL) was added dropwise and stirred for 10 mins before concentrating in vacuo. Flash chromatography purification as performed (silica, 2% → 5%MeOH in DCM) to yield a white solid (377 mg). 88% yield. (Supp. Data File 2).

**(*R*)-6-amino-7-(4-phenoxyphenyl)-9-(pyrrolidin-3-yl)-7,9-dihydro-8*H*-purin-8-one.HCl**

Boc-protected Tirabrutinib intermediate (*tert*-butyl (*R*)-3-(6-amino-8-oxo-7-(4-phenoxyphenyl)-7,8-dihydro-9*H*-purin-9-yl)pyrrolidine-1-carboxylate (AstaTech Inc., C92619), 500 mg, 1.0 mmoles) was dissolved in MeOH/ EtOAc (1:1, 30 mL), then a solution of HCl in dioxane (4M, 2.0 mL, 8 mmoles) was added and stirred at RT for 24 hrs. Volatiles were removed in vacuo, the resulting white solid triturated with hexane (3 x 5 mL) and dried to yield product as a white solid (374 mg). 86% Yield. (Supp. Data File 2).

**(*R*)-9-(1-acryloylpyrrolidin-3-yl)-6-amino-7-(4-phenoxyphenyl)-7,9-dihydro-8*H*-purin-8-one**

(*R*)-6-amino-7-(4-phenoxyphenyl)-9-(pyrrolidin-3-yl)-7,9-dihydro-8*H*-purin-8-one.HCl (180 mg, 0.42 mmoles) was suspended in DCM (10 mL) and cooled to 0 °C. DIPEA (221 µL, 164 mg, 1.27 mmoles) was added followed by acryloyl chloride (35 µL, 38 mg, 0.42 mmoles). The reaction mixture was stirred at 0 °C for 30 mins, then MeOH (1 mL) was added dropwise and stirred for 10 mins before concentrating in vacuo. Flash chromatography purification as performed (silica, 2% → 5%MeOH in DCM) to yield a white solid (145 mg). 78% yield. (Supp. Data File 2).

**(*R*)-9-(1-acetylpyrrolidin-3-yl)-6-amino-7-(4-phenoxyphenyl)-7,9-dihydro-8*H*-purin-8-one**

(*R*)-6-amino-7-(4-phenoxyphenyl)-9-(pyrrolidin-3-yl)-7,9-dihydro-8*H*-purin-8-one.HCl (180 mg, 0.42 mmoles) was suspended in DCM (10 mL) and cooled to 0 °C. DIPEA (221 μL, 164 mg, 1.27 mmoles) was added followed by acetyl chloride (30 μL, 33 mg, 0.42 mmoles). The reaction mixture was stirred at 0 °C for 30 mins, then MeOH (1 mL) was added dropwise and stirred for 10 mins before concentrating in vacuo. Flash chromatography purification as performed (silica, 2% → 5%MeOH in DCM) to yield a white solid (170 mg). 94% yield. (Supp. Data File 2).

References:


1. R. E. Joseph *et al.*, Impact of the clinically approved BTK inhibitors on the conformation of full-length BTK and analysis of the development of BTK resistance mutations in chronic lymphocytic leukemia. *Elife* **13** (2024).
2. Anonymous (The PyMOL Molecular Graphics System, Version 3.1.6.1 Schrödinger, LLC.
3. P. R. Evans, G. N. Murshudov, How good are my data and what is the resolution? *Acta Crystallogr D Biol Crystallogr* **69**, 1204-1214 (2013).
4. W. Kabsch, Xds. *Acta Crystallogr D Biol Crystallogr* **66**, 125-132 (2010).
5. C. Vonrhein *et al.*, Data processing and analysis with the autoPROC toolbox. *Acta Crystallogr D Biol Crystallogr* **67**, 293-302 (2011).
6. A. J. McCoy *et al.*, Phaser crystallographic software. *J Appl Crystallogr* **40**, 658-674 (2007).
7. O. S. Smart, Sharff A., Holstein, J., Womack, T.O., Flensburg, C., Keller, P., Paciorek, W., Vonrhein, C. and Bricogne G. (2021) (Cambridge, United Kingdom: Global Phasing Ltd.).
8. N. W. Moriarty, R. W. Grosse-Kunstleve, P. D. Adams, electronic Ligand Builder and Optimization Workbench (eLBOW): a tool for ligand coordinate and restraint generation. *Acta Crystallogr D Biol Crystallogr* **65**, 1074-1080 (2009).
9. P. Emsley, K. Cowtan, Coot: model-building tools for molecular graphics. *Acta Crystallogr D Biol Crystallogr* **60**, 2126-2132 (2004).
10. D. Liebschner *et al.*, Macromolecular structure determination using X-rays, neutrons and electrons: recent developments in Phenix. *Acta Crystallogr D Struct Biol* **75**, 861-877 (2019).
11. R. E. Joseph, T. E. Wales, D. B. Fulton, J. R. Engen, A. H. Andreotti, Achieving a Graded Immune Response: BTK Adopts a Range of Active/Inactive Conformations Dictated by Multiple Interdomain Contacts. *Structure* **25**, 1481-1494 e1484 (2017).
12. M. J. T. Frisch, G. W.; Schlegel, H. B.; Scuseria, G. E.; Robb, M. A.; Cheeseman, J. R.; Scalmani, G.; Barone, V.; Petersson, G. A.; Nakatsuji, H.; Li, X.; Caricato, M.; Marenich, A. V.; Bloino, J.; Janesko, B. G.; Gomperts, R.; Mennucci, B.; Hratchian, H. P.; Ortiz, J. V.; Izmaylov, A. F.; Sonnenberg, J. L.; Williams-Young, D.; Ding, F.; Lipparini, F.; Egidi, F.; Goings, J.; Peng, B.; Petrone, A.; Henderson, T.; Ranasinghe, D.; Zakrzewski, V. G.; Gao, J.; Rega, N.; Zheng, G.; Liang, W.; Hada, M.; Ehara, M.; Toyota, K.; Fukuda, R.; Hasegawa, J.; Ishida, M.; Nakajima, T.; Honda, Y.; Kitao, O.; Nakai, H.; Vreven, T.; Throssell, K.; Montgomery, J. A., Jr.; Peralta, J. E.; Ogliaro, F.; Bearpark, M. J.; Heyd, J. J.; Brothers, E. N.; Kudin, K. N.; Staroverov, V. N.; Keith, T. A.; Kobayashi, R.; Normand, J.; Raghavachari, K.; Rendell, A. P.; Burant, J. C.; Iyengar, S. S.; Tomasi, J.; Cossi, M.; Millam, J. M.; Klene, M.; Adamo, C.; Cammi, R.; Ochterski, J. W.; Martin, R. L.; Morokuma, K.; Farkas, O.; Foresman, J. B.; Fox, D. J., (2016) Gaussian 16, Revision C.01.
13. Y. Zhao, D. G. Truhlar, The M06 suite of density functionals for main group thermochemistry, thermochemical kinetics, noncovalent interactions, excited states, and transition elements: two new functionals and systematic testing of four M06-class functionals and 12 other functionals. *Theor Chem Acc* **120**, 215-241 (2008).

14. R. Valero, J. R. B. Gomes, D. G. Truhlar, F. Illas, Good performance of the M06 family of hybrid meta generalized gradient approximation density functionals on a difficult case: CO adsorption on MgO(001). *J Chem Phys* **129** (2008).
15. A. Lodola, D. Callegari, L. Scalvini, S. Rivara, M. Mor, Design and SAR Analysis of Covalent Inhibitors Driven by Hybrid QM/MM Simulations. *Methods Mol Biol* **2114**, 307-337 (2020).
16. N. Amatya *et al.*, Lipid-targeting pleckstrin homology domain turns its autoinhibitory face toward the TEC kinases. *Proc Natl Acad Sci U S A* **116**, 21539-21544 (2019).
17. R. E. Joseph *et al.*, Activation loop dynamics determine the different catalytic efficiencies of B cell- and T cell-specific tec kinases. *Sci Signal* **6**, ra76 (2013).
