## Supplemental Data File 2 for "BTK dynamics and function are differentially influenced by distinct covalent inhibitor warhead chemistries"

Alkane-Ibrutinib analog

**$^1$HNMR for RB5111 (400 MHz; DMSO-$d_6$)**

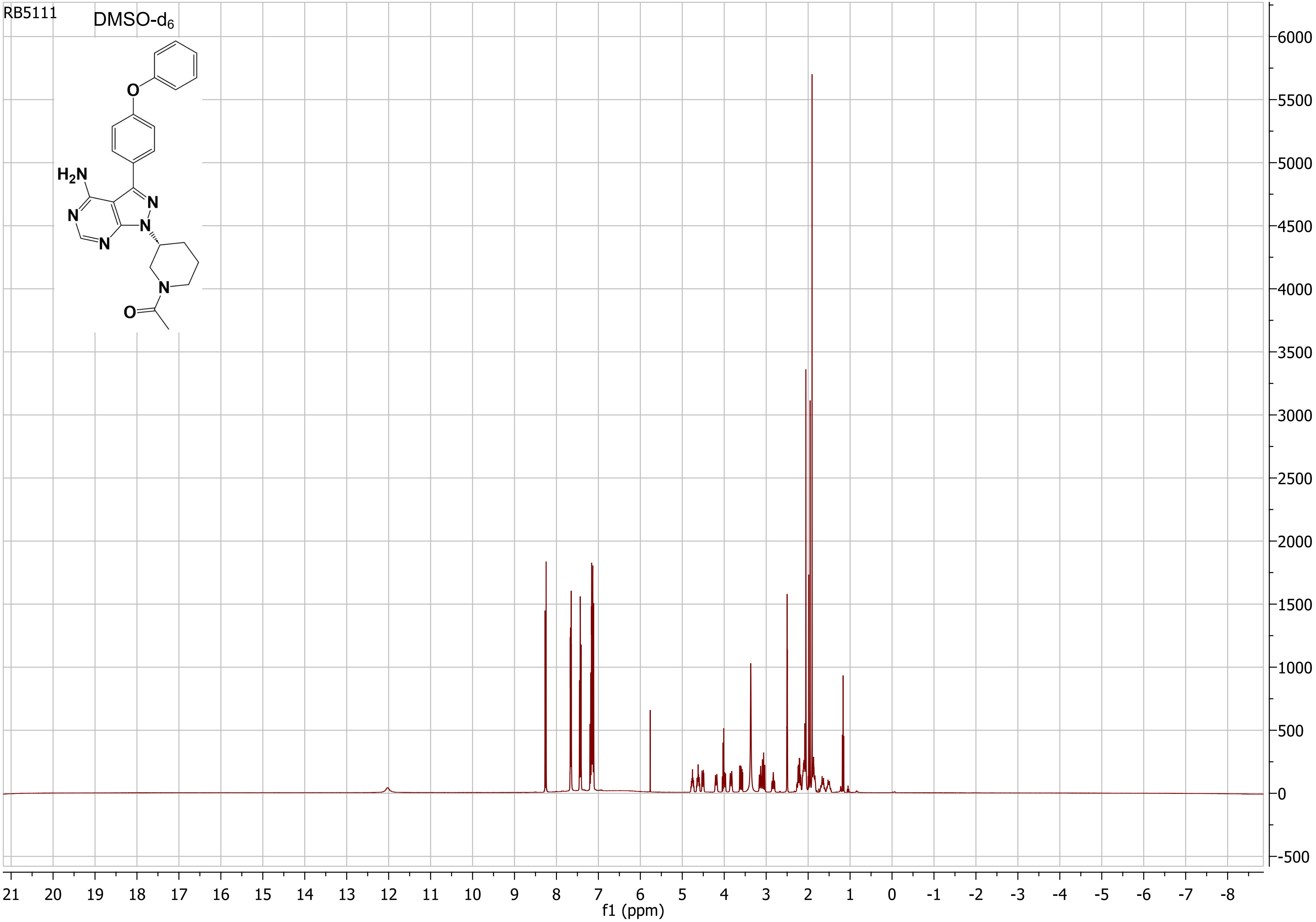

Alkane-Ibrutinib analog

#### $^{13}$CNMR for RB5111 (75 MHz; DMSO-$d_6$)

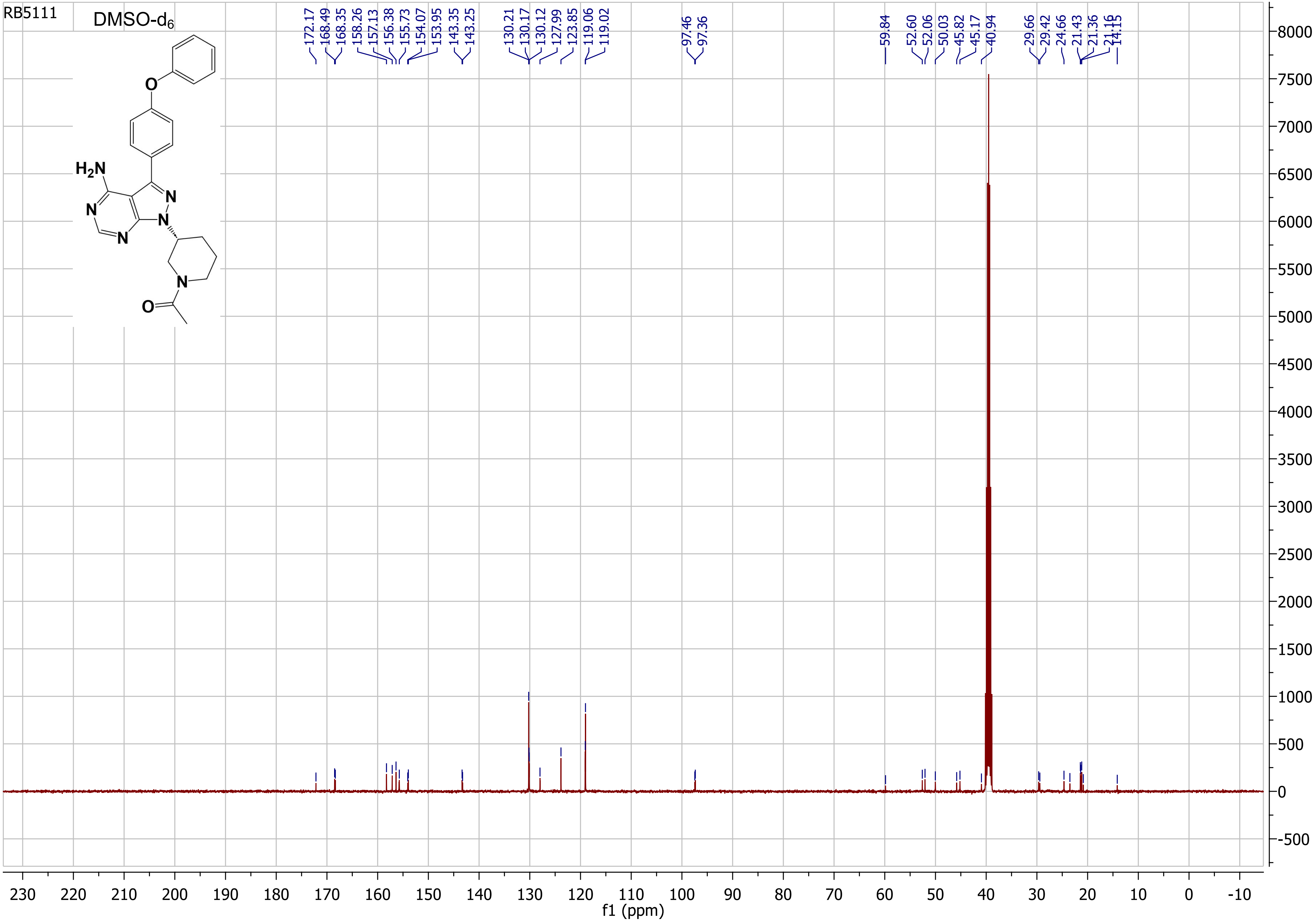

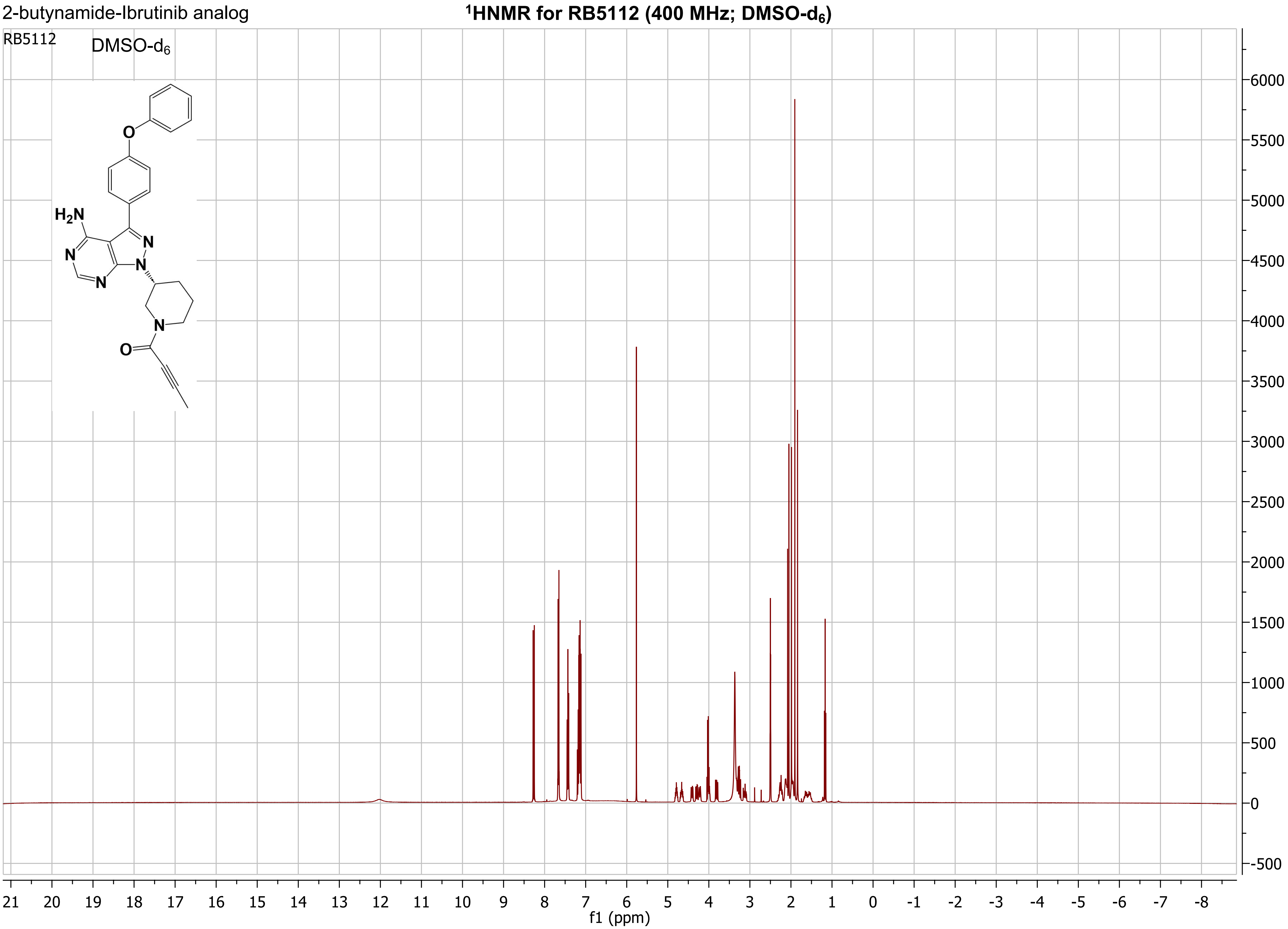
2-butynamide-Ibrutinib analog
¹HNMR for RB5112 (400 MHz; DMSO-d6)
RB5112
DMSO-d6
H2N
N
N
N
N
O
N
O
6000
5500
5000
4500
4000
3500
3000
2500
2000
1500
1000
500
0
-500
21 20 19 18 17 16 15 14 13 12 11 10 9 8 7 6 5 4 3 2 1 0 -1 -2 -3 -4 -5 -6 -7 -8
f1 (ppm)

2-butynamide-Ibrutinib analog

#### $^{13}$CNMR for RB5112 (75 MHz; DMSO-$d_6$)

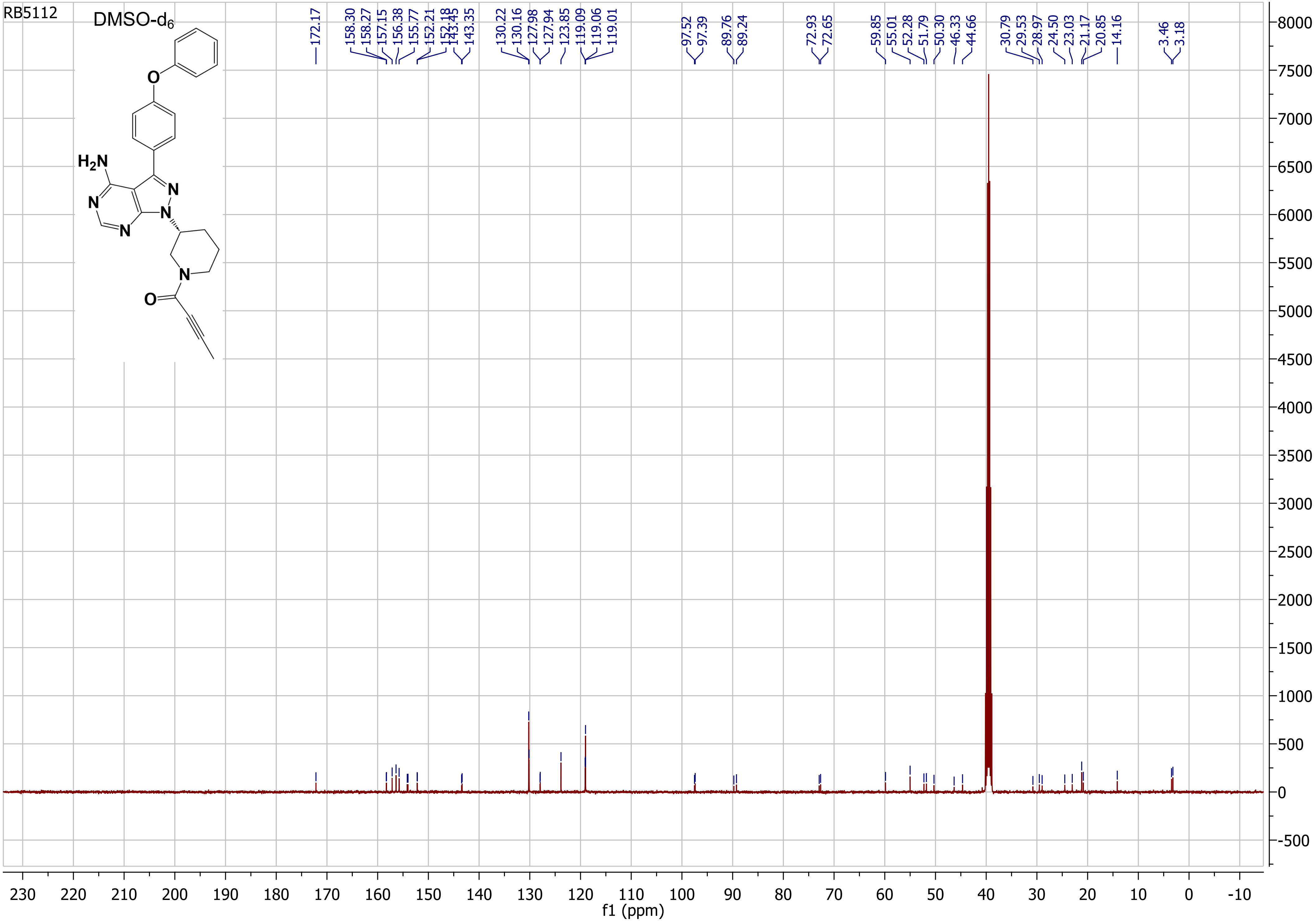

alkane-Tirabrutinib analog

### [1]HNMR for RB5123 (400 MHz; $CDCl_3$ + TMS)

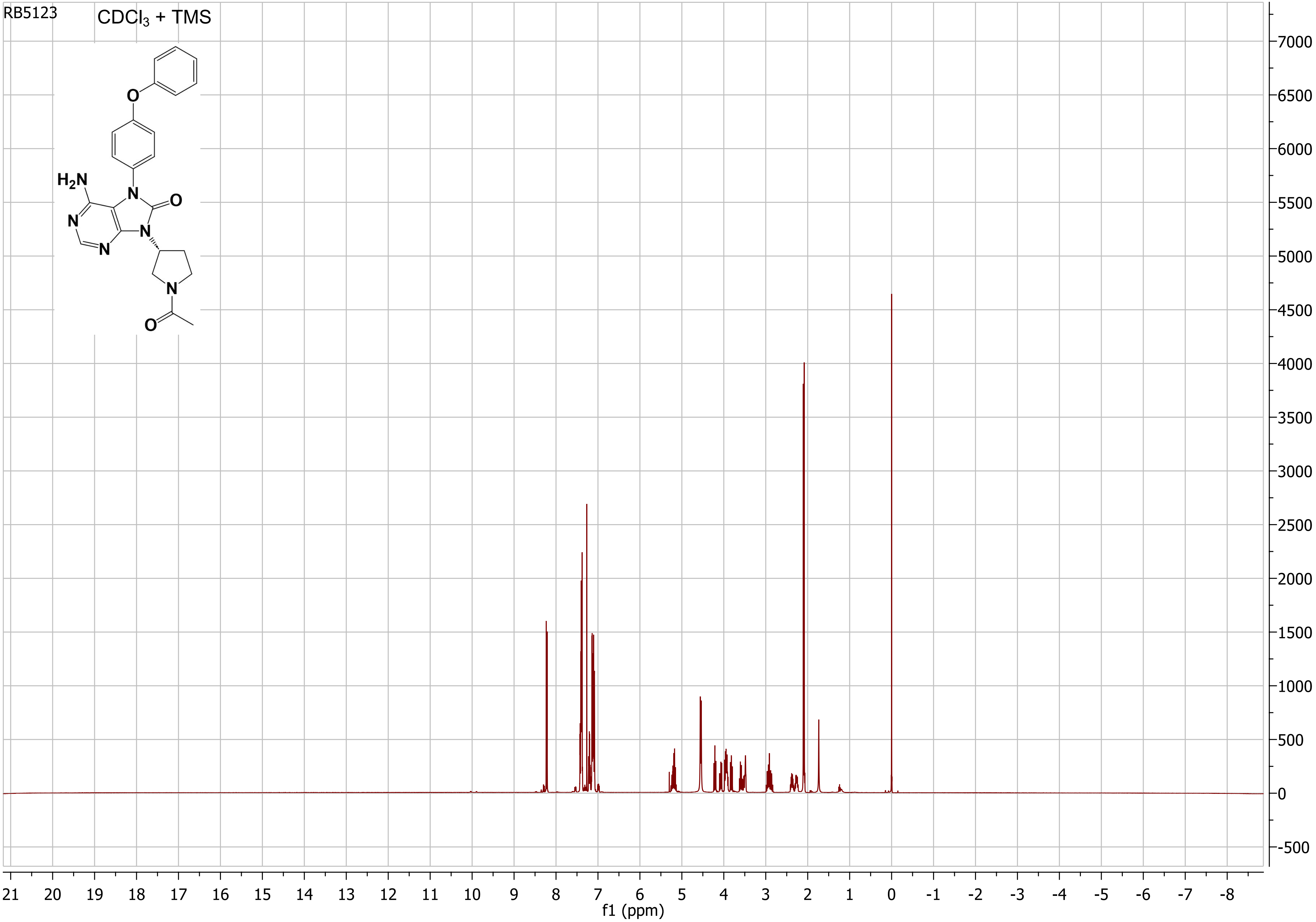

alkane-Tirabrutinib analog

**$^{13}$CNMR for RB5123 (75 MHz; $CDCl_3$ + TMS)**

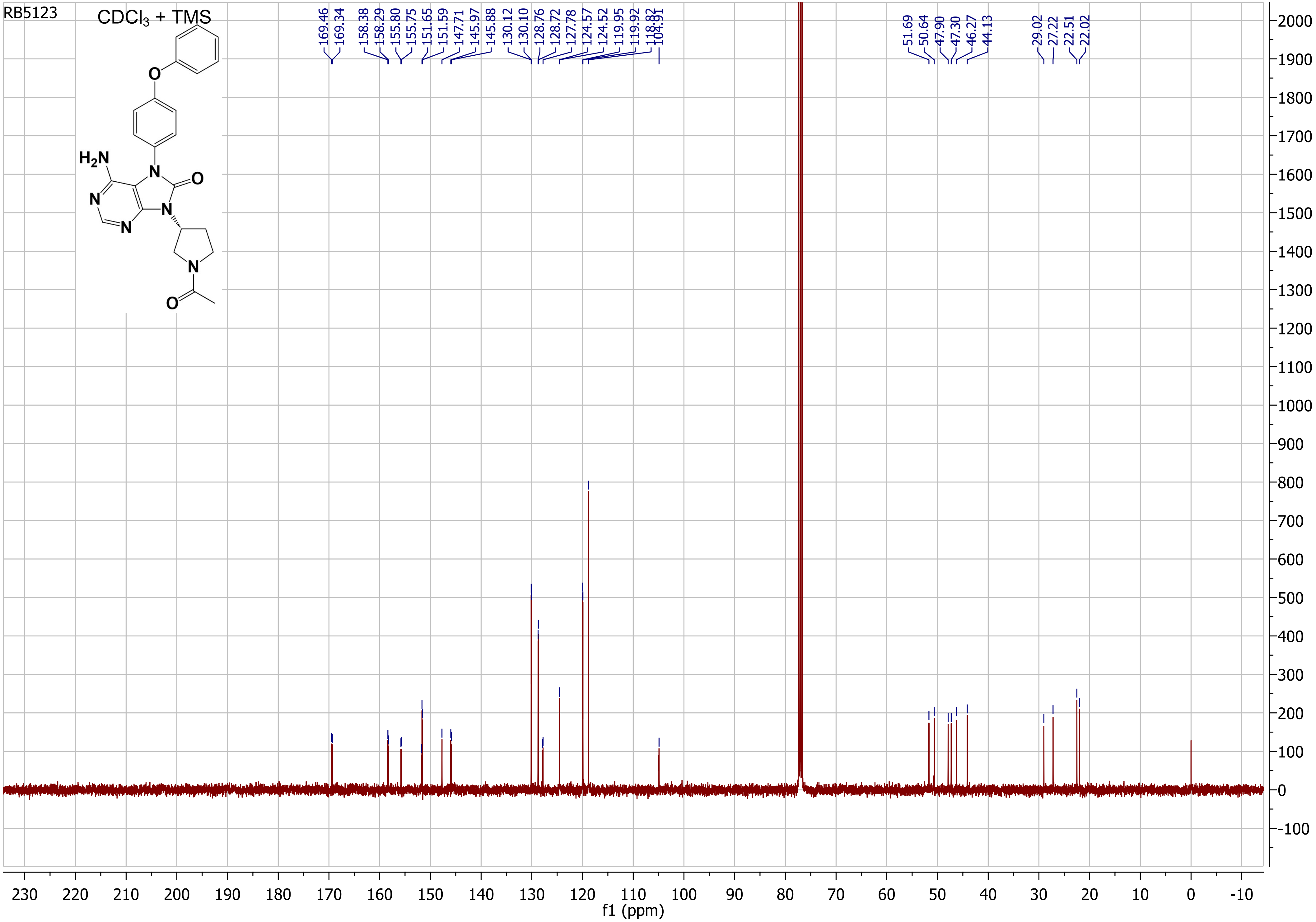

Acrylamide-Tirabrutinib analog

**$^{1}$HNMR for RB5124 (400 MHz; $CDCl_3$ + TMS)**

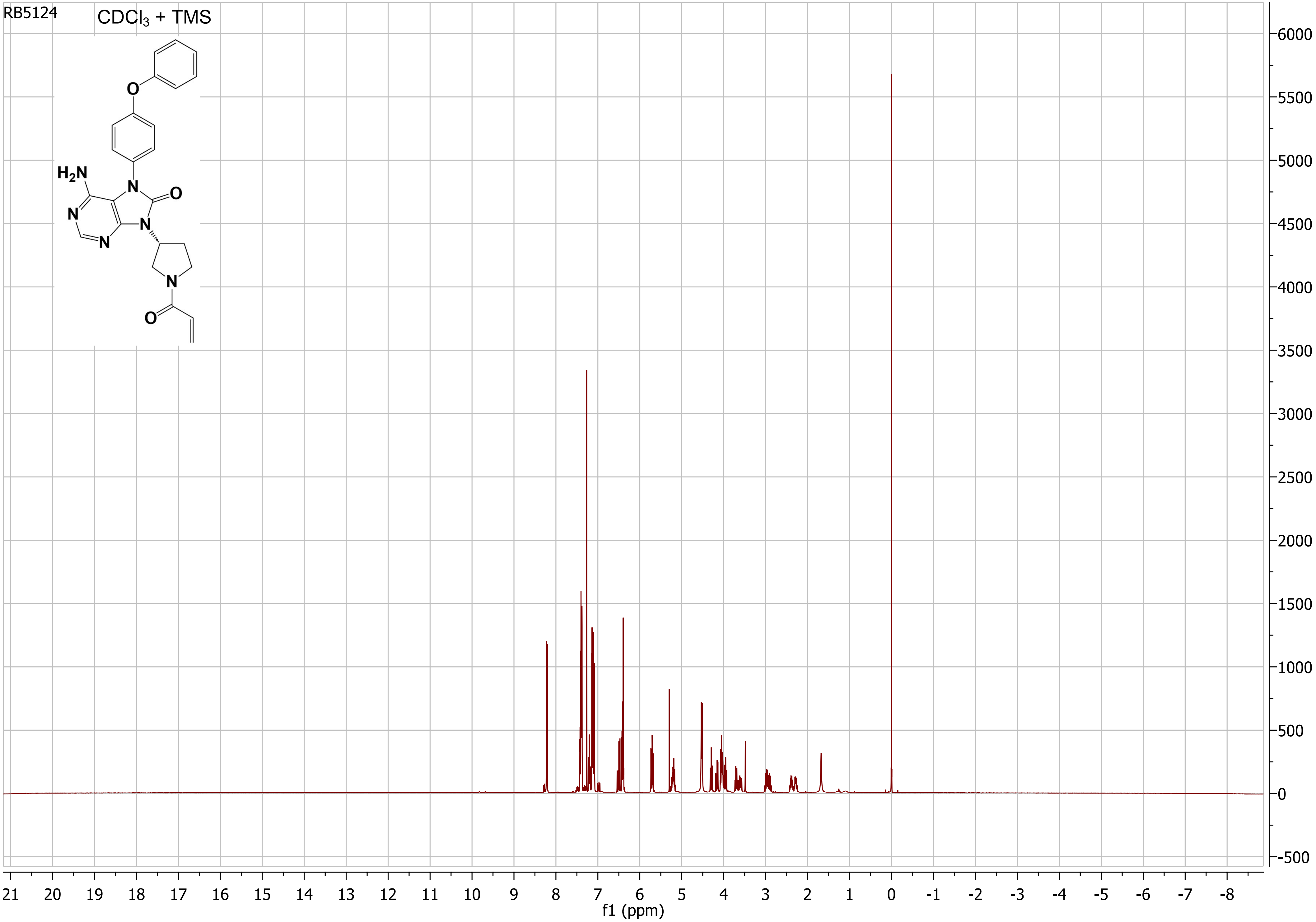

Acrylamide-Tirabrutinib analog

#### $^{13}$CNMR for RB5124 (75 MHz; $CDCl_3$ + TMS)

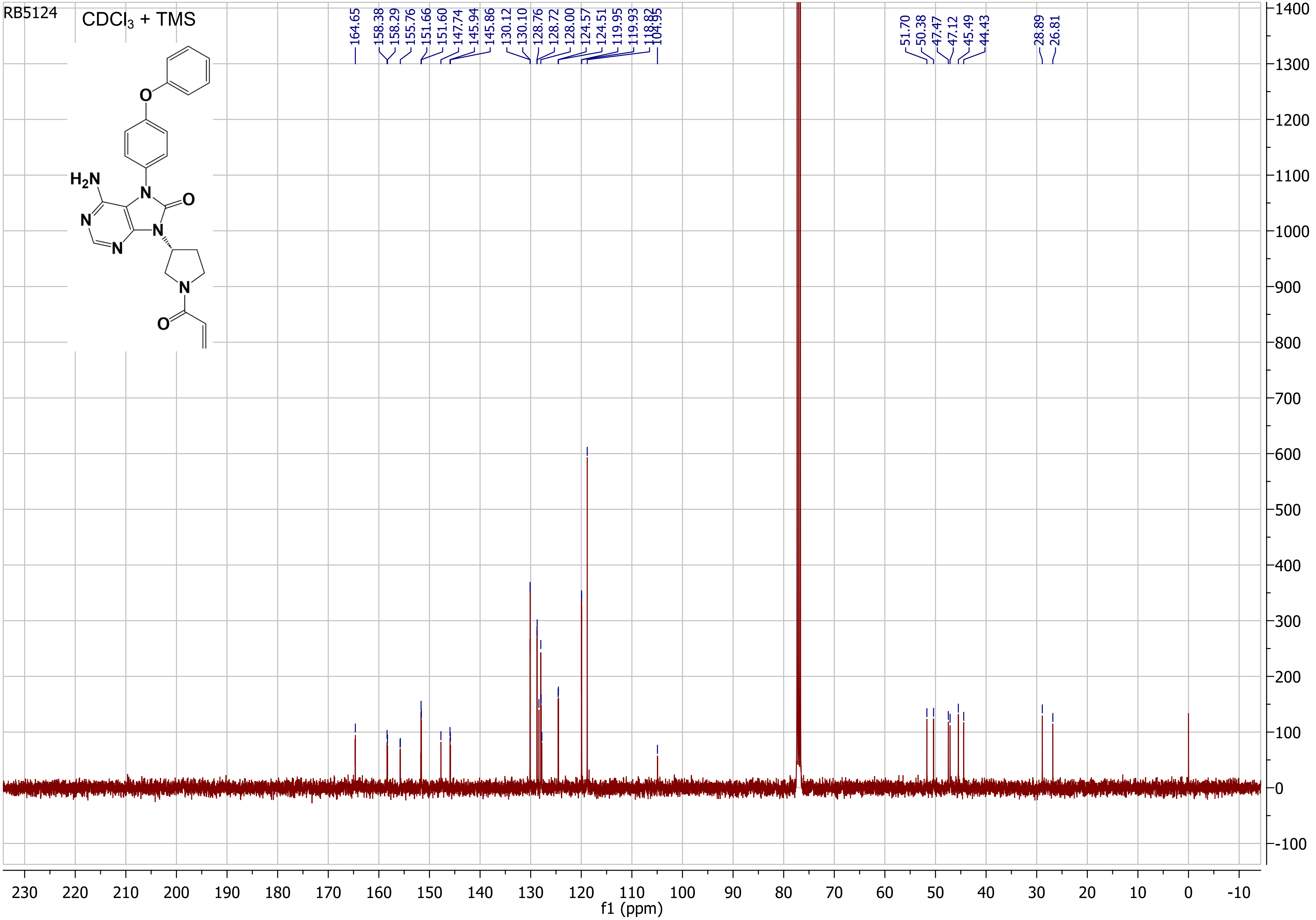

Mass spectrometry methods:

ESI LC-MS (Electrospray Ionisation Liquid Chromatography Mass Spectrometry)

The compounds were analysed by LC-MS using Xevo G2-XS QTof mass spectrometer (Waters) coupled to an Acquity LC system (Waters) using an Acquity UPLC BEH C18 column (2.1 x 50 mm, Waters). The flow rate was 0.6 ml min-1 and the gradient was as follows: 95% Solvent A (0.1% formic acid in water) with 5% solvent B (0.1% formic acid in acetonitrile) was held constant for 0.5 min, followed by a linear gradient to 100% solvent B over the next 2.1 min. After 1 min at 100% solvent B, the gradient was returned to 95% solvent A and 5% solvent B over 0.2 min and held for further 0.2min under this condition. The ESI capillary voltage was 2.5 kV, cone voltage 40 V and collision energy 6 eV. The MS acquisition rate was 10 spectra per second and m/z data ranging from 50 to 1200 Da was collected. Mass accuracy was achieved using a reference lock mass scan, once every 5 seconds. Data acquisition and processing was done using MassLynx v4.2 software (Waters).

**Accurate LC-MS (ESI) for RB511 showing:**
**A)** LC mobile-phase gradient (solvent A water (0.1% formic acid) and solvent B $CH_3CN$ (0.1% formic acid))
**B)** UV chromatogram response at 260 nm
**C)** Chromatogram set at 429 m/z
**D)** Base peak intensity (BPI) chromatogram showing most intense ion signal
**E)** Total ion chromatogram (TIC) showing the intensity of all detected ions

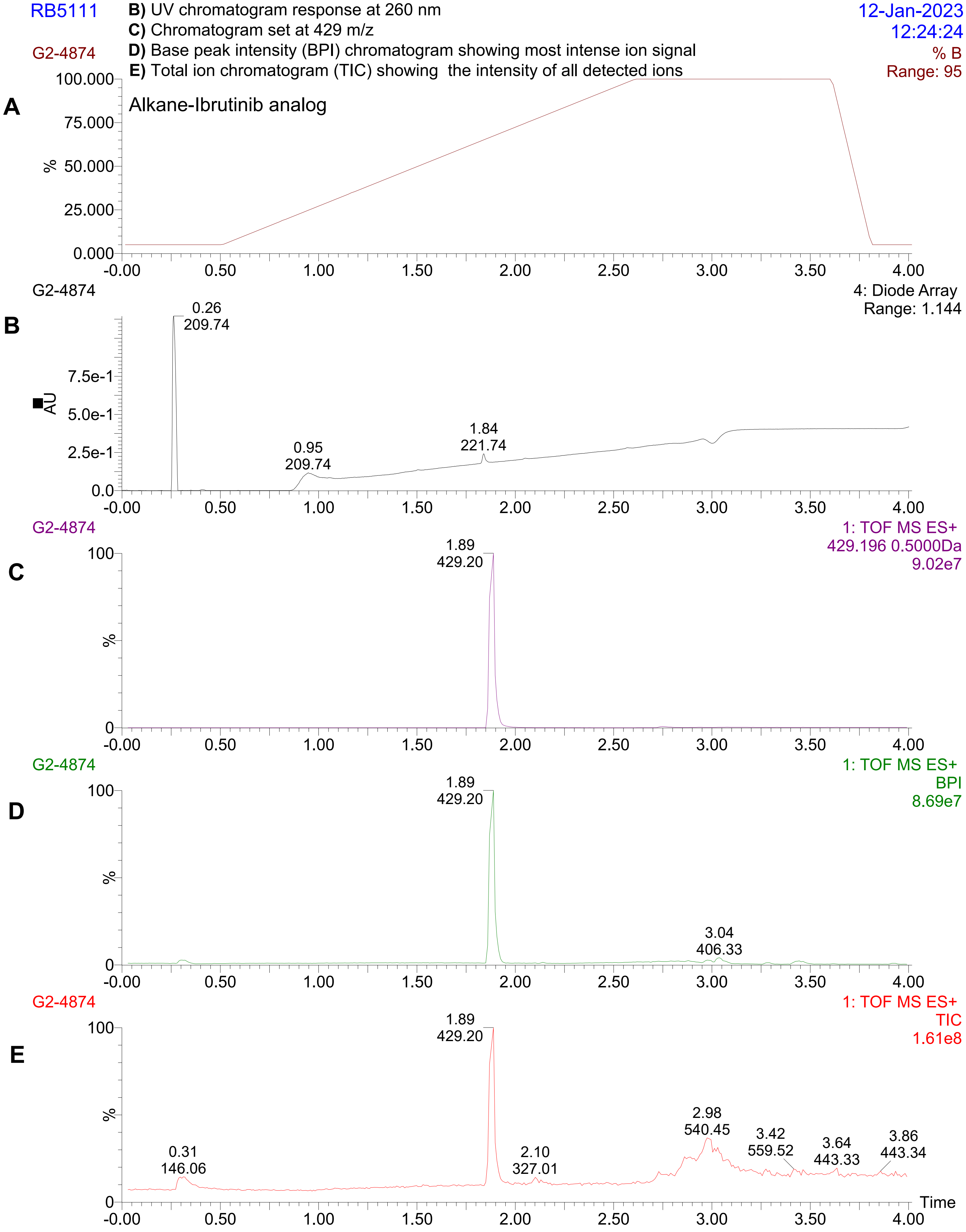

#### Accurate mass of RB5111 showing $[MH]^+$ and $[MNa]^+$ both within 5ppm of the calculated mass

**Elemental Composition Report** **$[MH]^+$**

Single Mass Analysis
Tolerance = 5.0 PPM / DBE: min = -1.5, max = 100.0
Element prediction: Off
Number of isotope peaks used for i-FIT = 3
Monoisotopic Mass, Even Electron Ions
437 formula(e) evaluated with 1 results within limits (up to 100 closest results for each mass)
Elements Used:
C: 24-24 H: 0-150 N: 0-30 O: 0-30

| | | | | | | | | |
|---|---|---|---|---|---|---|---|---|
| Minimum: | | | | -1.5 | | | | |
| Maximum: | | 5.0 | 5.0 | 100.0 | | | | |
| Mass | Calc. Mass | mDa | PPM | DBE | i-FIT | Norm | Conf(%) | Formula |
| 429.2040 | 429.2039 | 0.1 | 0.2 | 15.5 | 889.4 | n/a | n/a | C24 H25 N6 O2 |

**Elemental Composition Report** **$[MNa]^+$**

Single Mass Analysis
Tolerance = 5.0 PPM / DBE: min = -1.5, max = 100.0
Element prediction: Off
Number of isotope peaks used for i-FIT = 3
Monoisotopic Mass, Even Electron Ions
914 formula(e) evaluated with 1 results within limits (up to 100 closest results for each mass)
Elements Used:
C: 24-24 H: 0-150 N: 0-30 O: 0-30 Na: 0-1

| | | | | | | | | |
|---|---|---|---|---|---|---|---|---|
| Minimum: | | | | -1.5 | | | | |
| Maximum: | | 5.0 | 5.0 | 100.0 | | | | |
| Mass | Calc. Mass | mDa | PPM | DBE | i-FIT | Norm | Conf(%) | Formula |
| 451.1865 | 451.1858 | 0.7 | 1.6 | 15.5 | 409.4 | n/a | n/a | C24 H24 N6 O2 Na |

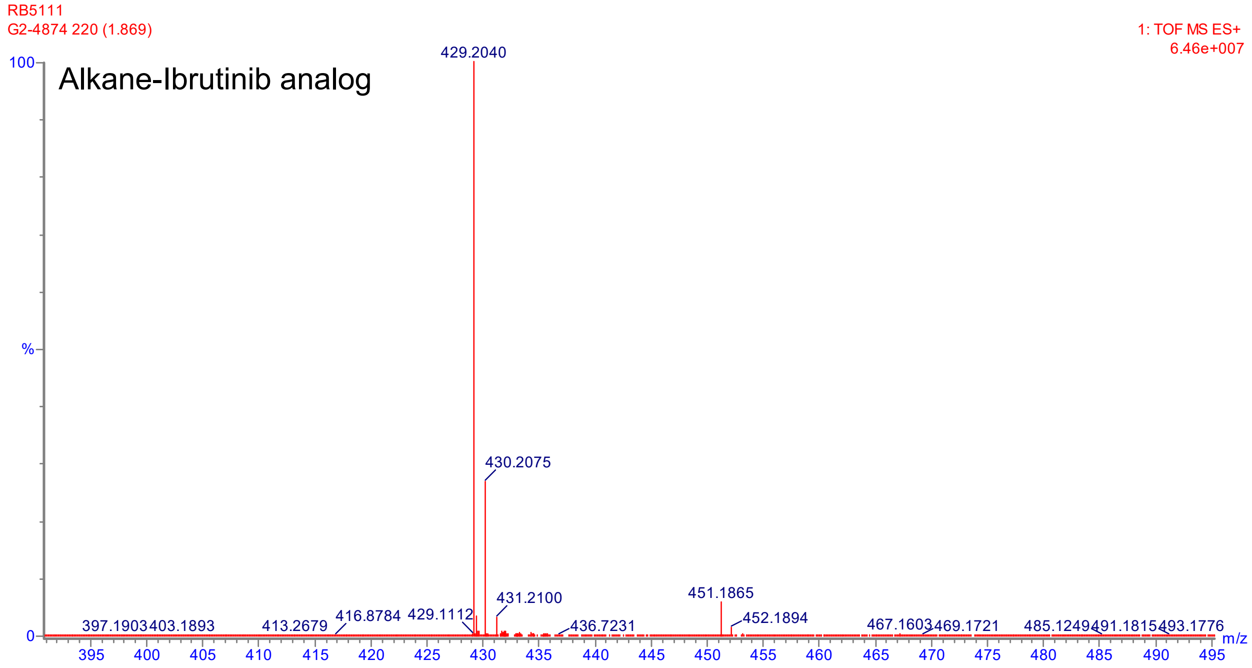

**Accurate LC-MS (ESI) for RB512 showing:**

**A)** LC mobile-phase gradient (solvent A water (0.1% formic acid) and solvent B $CH_3CN$ (0.1% formic acid))
**B)** UV chromatogram response at 260 nm
**C)** Chromatogram set at 429 m/z
**D)** Base peak intensity (BPI) chromatogram showing most intense ion signal
**E)** Total ion chromatogram (TIC) showing the intensity of all detected ions

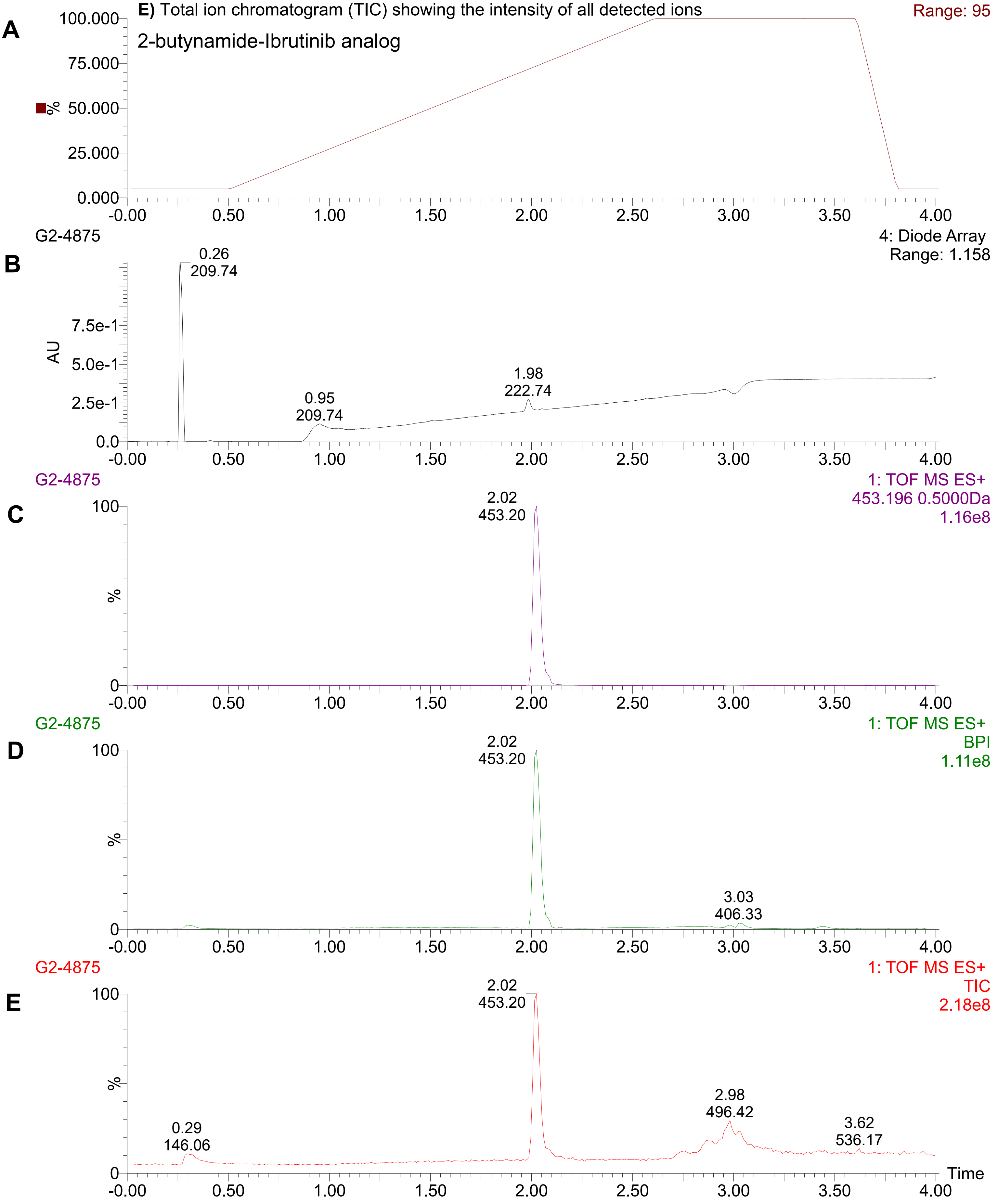

#### Accurate mass of RB5112 showing $[MH]^+$ and $[MNa]^+$ both within 5ppm of the calculated mass

**Elemental Composition Report** **$[MH]^+$**

Single Mass Analysis
Tolerance = 5.0 PPM / DBE: min = -1.5, max = 100.0
Element prediction: Off
Number of isotope peaks used for i-FIT = 3
Monoisotopic Mass, Even Electron Ions
483 formula(e) evaluated with 1 results within limits (up to 100 closest results for each mass)
Elements Used:
C: 26-26 H: 0-150 N: 0-30 O: 0-30

| | | | | | | | | |
|---|---|---|---|---|---|---|---|---|
| Minimum: | | | | -1.5 | | | | |
| Maximum: | | 5.0 | 5.0 | 100.0 | | | | |
| Mass | Calc. Mass | mDa | PPM | DBE | i-FIT | Norm | Conf(%) | Formula |
| 453.2045 | 453.2039 | 0.6 | 1.3 | 17.5 | 941.8 | n/a | n/a | C26 H25 N6 O2 |

**Elemental Composition Report** **$[MNa]^+$**

Single Mass Analysis
Tolerance = 5.0 PPM / DBE: min = -1.5, max = 100.0
Element prediction: Off
Number of isotope peaks used for i-FIT = 3
Monoisotopic Mass, Even Electron Ions
1007 formula(e) evaluated with 1 results within limits (up to 100 closest results for each mass)
Elements Used:
C: 26-26 H: 0-150 N: 0-30 O: 0-30 Na: 0-1

| | | | | | | | | |
|---|---|---|---|---|---|---|---|---|
| Minimum: | | | | -1.5 | | | | |
| Maximum: | | 5.0 | 5.0 | 100.0 | | | | |
| Mass | Calc. Mass | mDa | PPM | DBE | i-FIT | Norm | Conf(%) | Formula |
| 475.1857 | 475.1858 | -0.1 | -0.2 | 17.5 | 348.5 | n/a | n/a | C26 H24 N6 O2 Na |

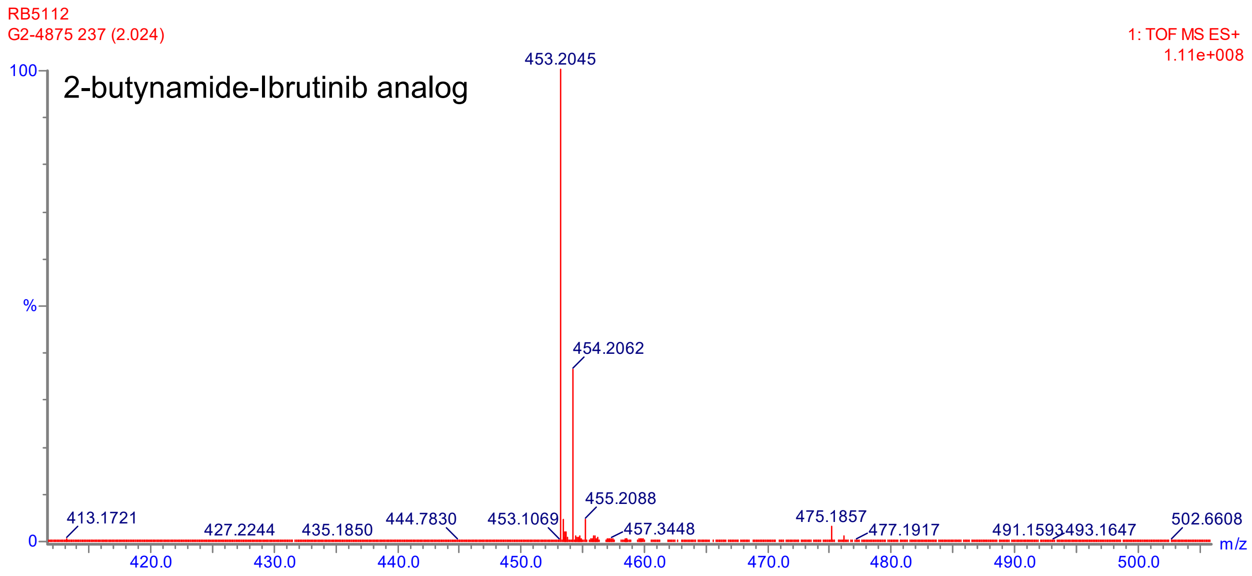

**Accurate LC-MS (ESI) for RB523 showing:**
**A)** LC mobile-phase gradient (solvent A water (0.1% formic acid) and solvent B $CH_3CN$ (0.1% formic acid))
**B)** UV chromatogram response at 260 nm
**C)** Chromatogram set at 429 m/z
**D)** Base peak intensity (BPI) chromatogram showing most intense ion signal
**E)** Total ion chromatogram (TIC) showing the intensity of all detected ions

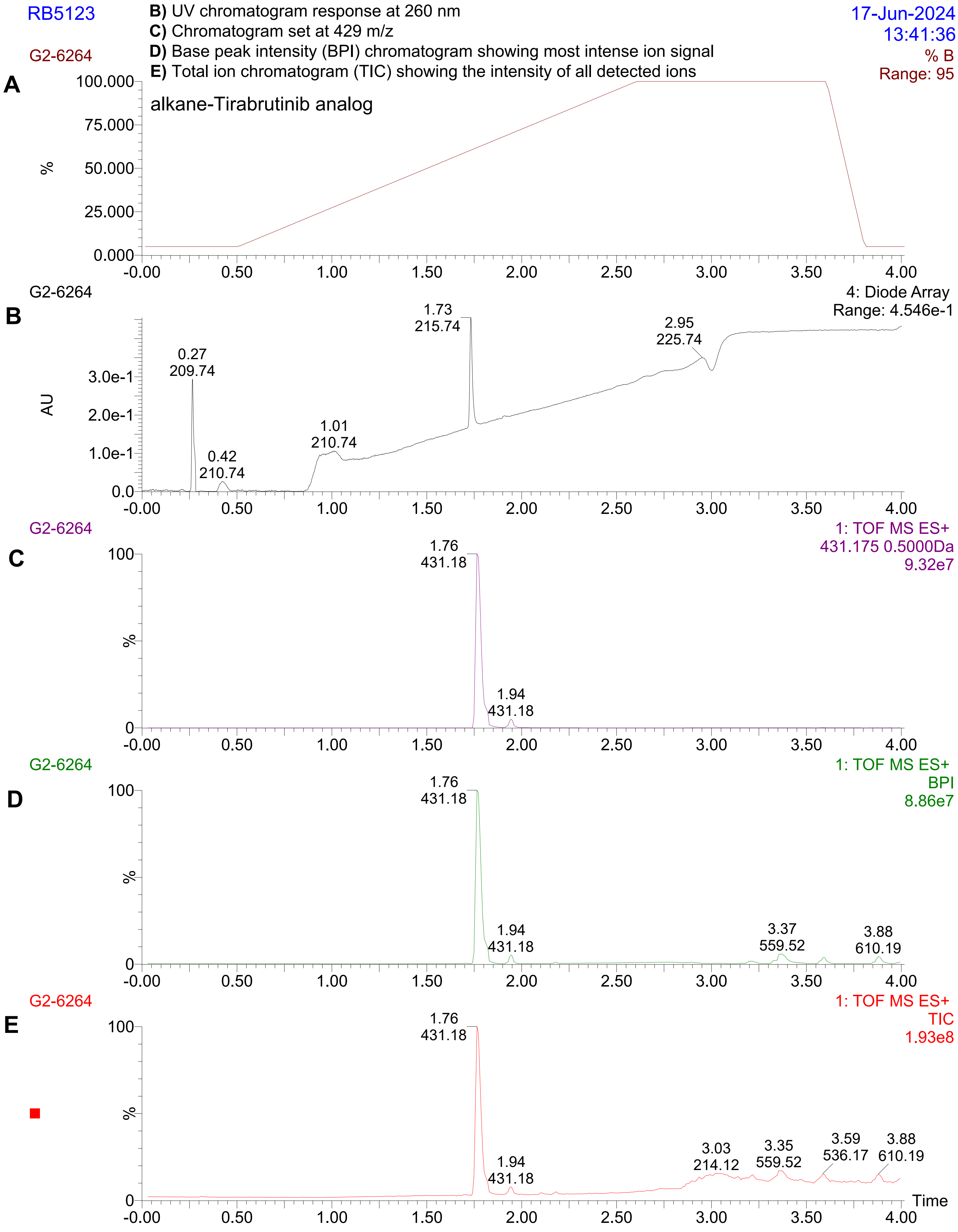

### Accurate mass of RB5123 showing [MH]$^+$ within 5ppm of the calculated mass

**Elemental Composition Report** **[MH]$^+$**

Single Mass Analysis
Tolerance = 5.0 PPM / DBE: min = -1.5, max = 100.0
Element prediction: Off
Number of isotope peaks used for i-FIT = 3

Monoisotopic Mass, Even Electron Ions
440 formula(e) evaluated with 1 results within limits (all results (up to 1000) for each mass)
Elements Used:
C: 23-23 H: 0-150 N: 0-30 O: 0-30

| | | | | | | | | |
|---|---|---|---|---|---|---|---|---|
| Minimum: | | | | -1.5 | | | | |
| Maximum: | | 5.0 | 5.0 | 100.0 | | | | |
| Mass | Calc. Mass | mDa | PPM | DBE | i-FIT | Norm | Conf(%) | Formula |
| 431.1835 | 431.1832 | 0.3 | 0.7 | 15.5 | 977.5 | n/a | n/a | C23 H23 N6 O3 |

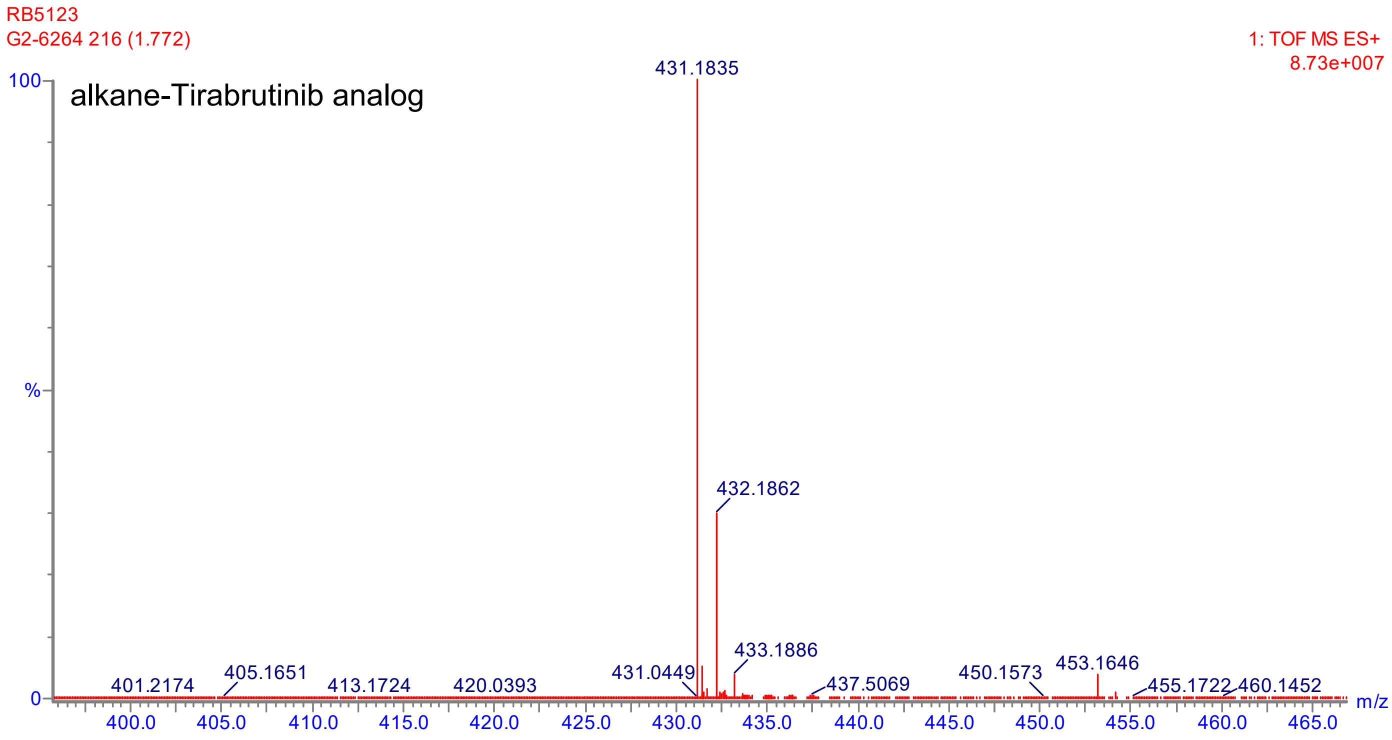

**Accurate LC-MS (ESI) for RB5124 showing:**
**A)** LC mobile-phase gradient (solvent A water (0.1% formic acid) and solvent B $CH_3CN$ (0.1% formic acid))
**B)** UV chromatogram response at 260 nm
**C)** Chromatogram set at 429 m/z
**D)** Base peak intensity (BPI) chromatogram showing most intense ion signal
**E)** Total ion chromatogram (TIC) showing the intensity of all detected ions

RB5124 — 18-Jun-2024 13:54:48

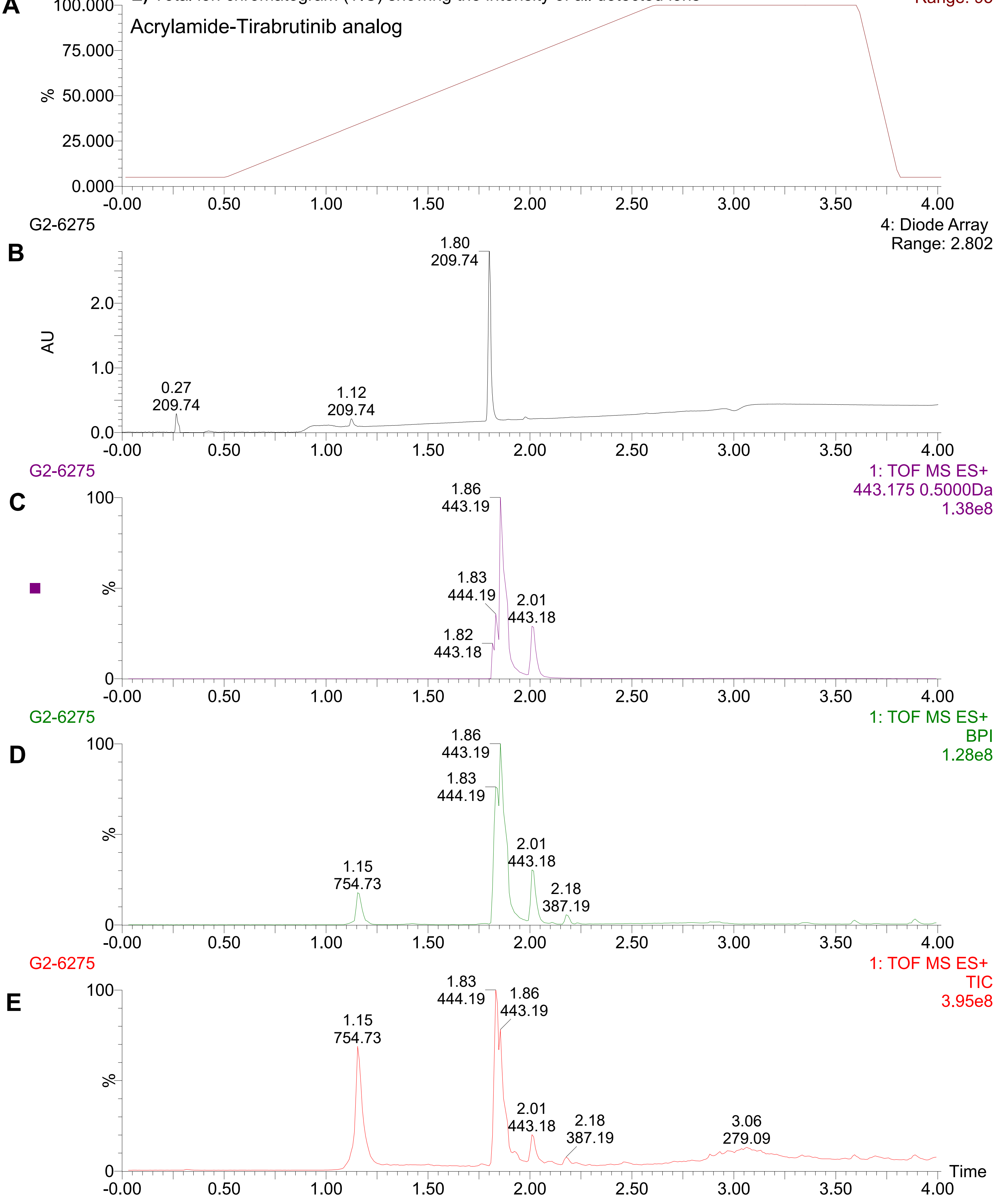

#### Accurate mass of RB5124 showing [MH]$^+$ within 5ppm of the calculated mass

**Elemental Composition Report [MH]$^+$**

Single Mass Analysis
Tolerance = 5.0 PPM / DBE: min = -1.5, max = 100.0
Element prediction: Off
Number of isotope peaks used for i-FIT = 3

Monoisotopic Mass, Even Electron Ions
463 formula(e) evaluated with 1 results within limits (all results (up to 1000) for each mass)
Elements Used:
C: 24-24 H: 0-150 N: 0-30 O: 0-30

| | | | | | | | | |
|---|---|---|---|---|---|---|---|---|
| Minimum: | | | | -1.5 | | | | |
| Maximum: | | 5.0 | 5.0 | 100.0 | | | | |
| Mass | Calc. Mass | mDa | PPM | DBE | i-FIT | Norm | Conf(%) | Formula |
| 443.1834 | 443.1832 | 0.2 | 0.5 | 16.5 | 981.3 | n/a | n/a | C24 H23 N6 O3 |

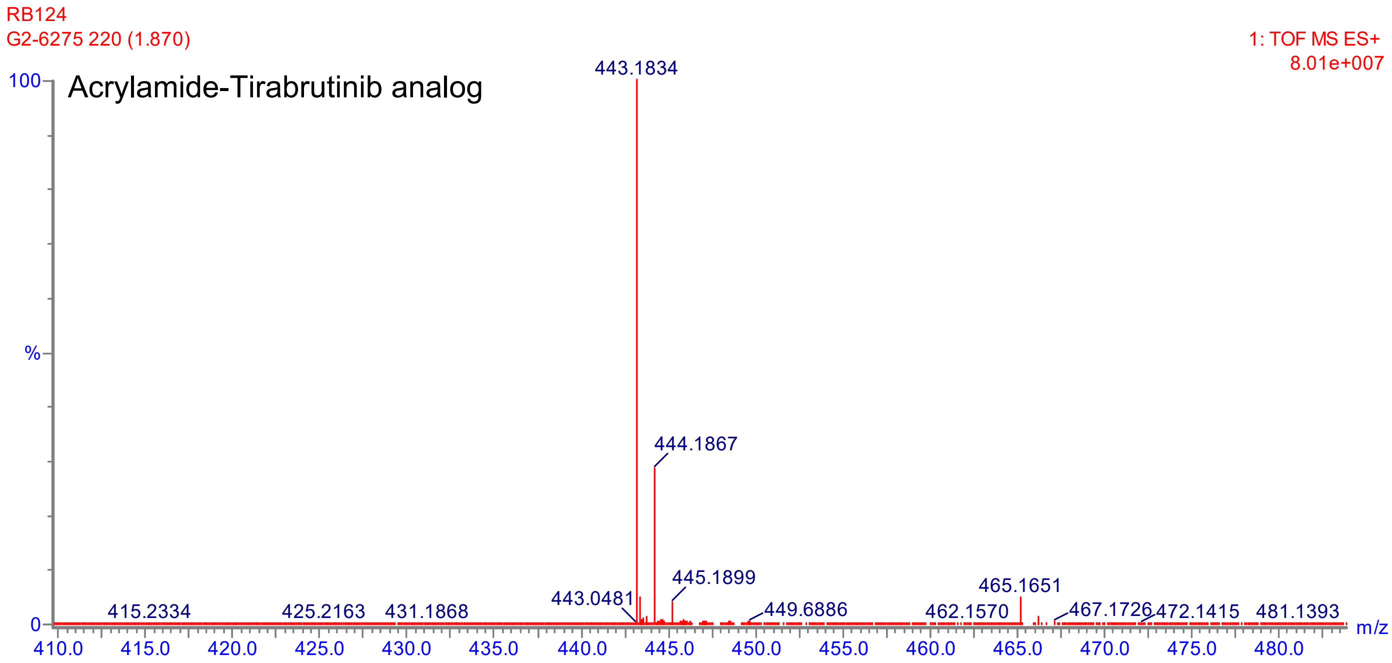
